# Host cholesterol dependent activation of VapC12 toxin enriches persister population during *Mycobacterium tuberculosis* infection

**DOI:** 10.1101/856286

**Authors:** Sakshi Talwar, Manitosh Pandey, Chandresh Sharma, Rintu Kutum, Josephine Lum, Daniel Carbajo, Renu Goel, Michael Poidinger, Debasis Dash, Amit Singhal, Amit Kumar Pandey

## Abstract

A worldwide increase in the frequency of multidrug-resistant and extensively drug-resistant cases of tuberculosis is mainly due to therapeutic noncompliance associated with a lengthy treatment regimen. Depending on the drug susceptibility profile, the treatment duration can extend from 6 months to 2 years. This protracted regimen is attributed to a supposedly non-replicating and metabolically inert subset of the *Mycobacterium tuberculosis* (Mtb) population, called ‘persisters’. The mechanism underlying stochastic generation and enrichment of persisters is not fully known. We have previously reported that the utilization of host cholesterol is essential for mycobacterial persistence. In this study, we have demonstrated that cholesterol-induced activation of a ribonuclease toxin (VapC12) inhibits translation by targeting proT tRNA in Mtb. This results in cholesterol-specific growth modulation that increases the frequency of the generation of persisters in a heterogeneous Mtb population. Also, a null mutant strain of this toxin (Δ*vapC12*) failed to persist and demonstrated an enhanced growth phenotype in a guinea pig model of Mtb infection. Thus, we have identified a novel strategy through which cholesterol-specific activation of a toxin–antitoxin (TA) module in Mtb enhances persister formation during infection. In addition to identifying the mechanism, the study provides opportunity for targeting persisters, a new paradigm facilitating tuberculosis drug development.

## Introduction

Globally, a third of the human population is infected with *Mycobacterium tuberculosis* (Mtb), the causative agent of tuberculosis. Being an obligate intracellular pathogen, Mtb has co-evolved with humans for centuries (*2–4*). Unlike the actively infected population, the latently infected individuals harbour Mtb for decades without showing any overt symptoms. This phenotype of Mtb is attributed to a slow-growing, metabolically altered subset of the heterogeneous Mtb population called persisters (*5, 6*). These persisters are refractory to antimycobacterial drugs and can only be targeted using a strict regimen consisting of a combination of drugs for an unusually extended period. The protracted regimen triggers noncompliance and results in an increased frequency of multidrug-resistant (MDR) and extensively drug-resistant (XDR) tuberculosis cases(*7–10*).

Persisters are extremely drug tolerant sub-population that possess an extraordinary ability to hide within a host. Although several studies have described stress-induced generation of persisters under in-vitro growth conditions (*11–14*), the exact conditions triggering the generation and enrichment of persisters inside the host during a normal course of mycobacterial infection remain unclear. Upon infection Mtb induces the formation of lipid-rich foamy macrophages. Lysis of these macrophages results in the formation of the caseous core of a typical ‘tuberculous granuloma’, providing Mtb with a cholesterol-rich niche. While residing in this nutrient-deprived granuloma, Mtb adapts itself to utilize cholesterol as a favoured carbon source (*15*). This cholesterol utilization causes inhibition of growth and activation of pathways leading to the generation of persisters in the mycobacterial population (*15, 16*). Mtb facilitate intracellular accumulation of cholesterol by up regulating cholesterol biosynthesis pathways that convert resident macrophages into foamy macrophages (*17*). These findings imply that Mtb hijacks host pathways to build a favourable niche for itself in order to remain as a ‘persister’ for decades, facilitating long-term persistence, which is a hallmark of mycobacterial pathogenesis (*18, 19*).

Toxin–antitoxin (TA) proteins play a crucial role in generating persisters in several bacterial species(*20–22*). These TA systems, known to modulate growth under various growth and stress conditions, are found in wide range of bacterial and archaeal chromosomes and plasmids (*23–25*). Research conducted during the past decade has clearly demonstrated that TA loci act as effectors of dormancy and persistence in several bacterial species (*20, 21*). Each TA locus consists of genes expressing a pair of toxin-antitoxin protein. Antitoxin, being more labile, degrades under specific growth and stress conditions resulting in the activation of cognate toxin. The activated toxin modulates growth by targeting growth related genes.

The genome of Mtb constitute 88 TA systems whereas saprophytic soil dwelling *Mycobacterium smegmatis* genome has only one TA locus, clearly highlighting the role of TA systems in bacterial adaptation and survival in a very hostile environment inside the host (*26*). Based on the mechanism of toxin activation, the TA system is classified into seven different types. The most characterized of all, the VapBC family, belongs to type II group. The toxin from the type II group targets all forms of RNA including mRNA, rRNA and tRNA. Although, the type II TA system has been shown to regulate persistence in several bacterial species, the exact mechanism is not very clear. It is hypothesized that each TA pair is required for survival of bacteria under specific growth and stress condition (*27, 28*), however presence of a very high number of the TA system in Mtb genome also increases the chances of redundancy and the possibility of multiple TA systems regulating specific growth conditions.

In the current study, we have identified the role of one such Mtb ribonuclease toxin, VapC12, to be critical for cholesterol-induced generation of antibiotic persistence in mycobacteria. Our data conclusively demonstrate that cholesterol activates the ribonuclease toxin by disrupting its binding to the cognate anti-toxin VapB12. We further demonstrated that the proT tRNA of Mtb is a bonafide substrate of the VapC12 ribonuclease toxin and that the toxin mediated modulation of the proT tRNA regulate the generation and enrichment of the cholesterol-induced persisters in mycobacteria. Finally, we also demonstrated that the *vapC12* dependent enrichment of antibiotic persistence also contribute towards disease persistence as seen in a guinea pig model of tuberculosis infection.

## Results

### *vapC12* gene is essential for cholesterol-specific growth modulation in Mycobacterium tuberculosis *(Mtb)*

We have previously demonstrated that utilization of host cholesterol as a carbon source is essential for maintaining persistence during Mtb infection(*15*). In this study, we have used Mtb grown in a cholesterol-rich media as an *in-vitro* model to examine the role of cholesterol in the formation, maintenance, and enrichment of persisters during Mtb infection. We have initially analysed the metabolic and replication rates of Mtb grown in the cholesterol-rich media. We have observed a decrease in the replication and metabolic rates of wild-type (WT) H37Rv grown in the cholesterol-rich media by ten- and three-fold, respectively, compared with (WT) H37Rv grown in the glycerol-rich media (Fig. 1A, 1B). An *in-vitro* time-kill curve assay revealed a cholesterol-specific increase in the frequency of the generation of a rifamycin-tolerant sub-population (Fig. 1C). The TA loci across bacterial species regulate growth(*21, 29–31*), therefore we speculated the aforementioned phenotype to be regulated by one of the Mtb TA loci. For that we analysed the data of a transposon mutagenesis screening performed in Mtb H37Rv to identify genes essential for growth in a cholesterol-rich environment(*32*). Through manual curation of data, we identified transposon insertions in six VapC toxins that were over-represented in cholesterol than in glycerol. This finding suggests the role of one or all of these toxins in the cholesterol-mediated growth modulation of Mtb (Fig. S1). Of these six *vapC* genes, we generated clean deletion mutants for the top two toxins VapC8 and VapC12, which demonstrated the highest increase in the growth rate. We found that compared with the WT strain, the *vapC8*-null strain showed no significant growth difference in the cholesterol-rich media (Fig. S2), whereas the mutant lacking *vapC12* gene failed to slow down its growth and was metabolically more active in the cholesterol-rich media, suggesting essentiality of this gene in cholesterol-mediated growth retardation. The mutant phenotype was found to be gene specific because adding back the *vapBC12* locus restored the WT phenotype (Fig. 1D, 1E). As expected, the cholesterol-mediated increase in the frequency of the generation of the rifamycin-tolerant population, observed in the WT strain, was reversed in the *vapC12*-null strain, underscoring the role of this putative toxin in the generation of cholesterol-induced persistence in mycobacteria (Fig. 1F, S3). To gain additional insights, we performed transcriptional profiling of these strains in both glycerol only and cholesterol only media through RNAseq analysis. As expected, the transcript levels of genes involved in cholesterol metabolism, methyl-citrate cycle, and glyoxylate pathways were significantly upregulated(*33, 34*). The slowing down of WT Mtb in the cholesterol-rich media can be attributed to a decrease in the transcript levels of genes involved in respiration (e.g. cytochrome and ATP synthesis pathway genes). A significant decrease in the expression of genes belonging to the *esx3* locus in WT Mtb was intriguing, suggesting the possibility of iron-mediated growth modulation in the cholesterol-rich environment(*35, 36*). Additionally we also observed an increase in the transcript levels of DosR regulon genes in the cholesterol-rich media (Fig. 1G, S4, Table S1). Despite the differences observed in the growth phenotype between H37Rv and *vapC12* mutants grown in the cholesterol-rich media, expression profiling data showed few differentially expressed genes between the samples, suggesting that a post-transcriptional regulation mechanism plays a decisive role in inducing cholesterol-mediated growth regulation in Mtb (Table S2). Since ATP depletion is one of the mechanisms through which bacteria increase their tolerance to antibiotics leading to persistence (*20, 37, 38*) and our RNA sequencing data also revealed differential expression of ATP synthesis pathway genes, we quantified intracellular ATP levels in glycerol- and cholesterol-grown BCG cultures. Compared with the glycerol-grown BCG culture, the cholesterol-grown BCG culture demonstrated a 25-fold decrease in intracellular ATP levels. This cholesterol-specific depletion of intracellular ATP depends on the presence of *vapC12* gene (Fig. 1H).

**Figure 1:**
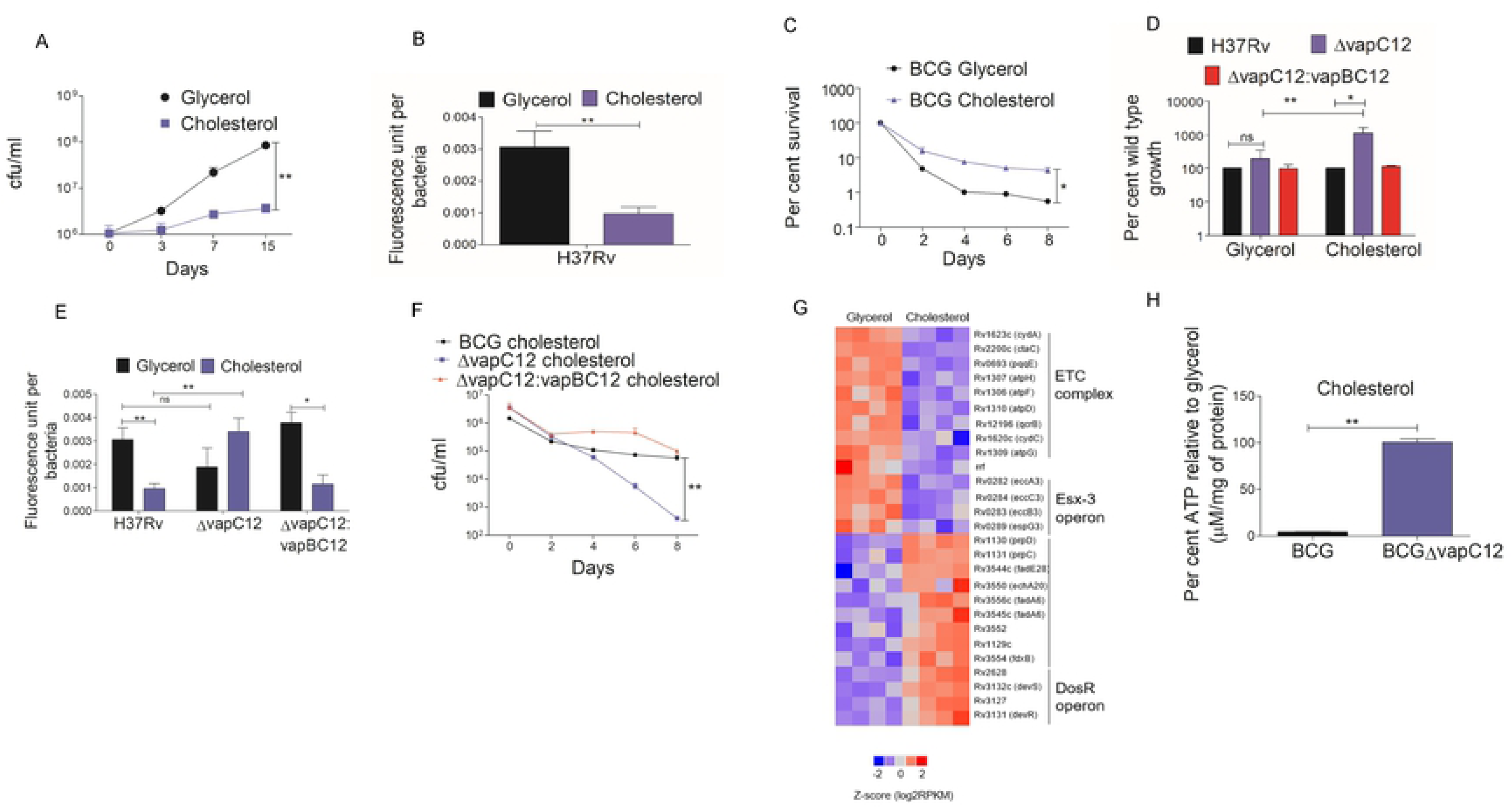
*vapC12* gene is essential for cholesterol-specific growth modulation in Mycobacterium tuberculosis *(Mtb)* A) The growth curve of H37Rv in a minimal media supplemented with 0.1% glycerol and 0.01% cholesterol. The log-phase cultures of H37Rv grown in 7H9 media enriched with OADC were washed with PBS-tyloxapol and resuspended in respective media at an absorbance of 0.005. Growth was estimated by CFU plating on 7H11+OADC plates at different time points post inoculation. Experiments were performed in triplicates, and data represent the mean ± SEM. B&E) Resazurin-based estimation of the metabolic activity of H37Rv (B) and *ΔvapC12* and *ΔvapC12:vapBC12* (E) grown in a minimal media supplemented with glycerol and cholesterol. Strains were serially diluted in a 96-well plate in respective media. The experiment was performed in two independent sets, and the plate was incubated at 37°C for 5 days. One set of the experiment was used for recording fluorescence after adding the presto blue reagent at 570 or 585 nm, whereas the other set was used for enumeration of bacteria present in each well. The metabolic activity calculated for each well is representative of the mean fluorescent readout per bacteria from three independent experiments. Data were analysed using unpaired Student’s t test. *P < 0.05, **P < 0.01 C&F) Kill curve of *M. bovis* BCG (C) and BCGΔ*vapC12* and BCG Δ*vapC12*:*vapBC12* (F) grown in glycerol- and cholesterol-rich media. Log-phase cultures of strains were washed with PBS-tyloxapol and inoculated at an absorbance of 0.05. The cultures were allowed to grow for 4 days before being treated with 5× MIC of rifamycin. Bacterial enumeration was performed through CFU plating of cultures on 7H11+OADC plates at various time points. The kill curve was plotted by calculating the percent survival. The experiment was repeated three times, and data represented are the mean ± SEM. Data were analysed using unpaired Student’s t test. *P < 0.05, **P < 0.01 D) Percent wild-type growth of the *vapC12* mutant and *ΔvapC12:vapBC12* in a minimal media containing 0.1% glycerol and 0.01% cholesterol. Growth was estimated by CFU plating of cultures on 7H11+OADC plates 8 days post inoculation. The experiment was repeated three times, and data represented are the mean ± SEM. Data were analysed using unpaired Student’s t test. *P < 0.05, **P < 0.01 G) Heat-map visualization of differentially expressed transcripts in wild type H37Rv grown in Glycerol and Cholesterol media, analysed through RNA-sequencing. Expression data of the respective genes based on FDR adjusted are depicted in the heat map. The RNA for sequencing was isolated from four different set of cultures grown in respective media. H) Percent estimation of ATP in wild-type BCG and *ΔBCGvapC12* strains grown in a cholesterol-rich media relative to glycerol-rich media. ATP estimated in micromolar concentrations was normalized with per milligram of protein in each sample. The experiment was repeated three times, and data plotted represents the mean ± SEM. Data were analysed using unpaired Student’s t test. P value *<0.05, **<0.01.

**Figure 2:**
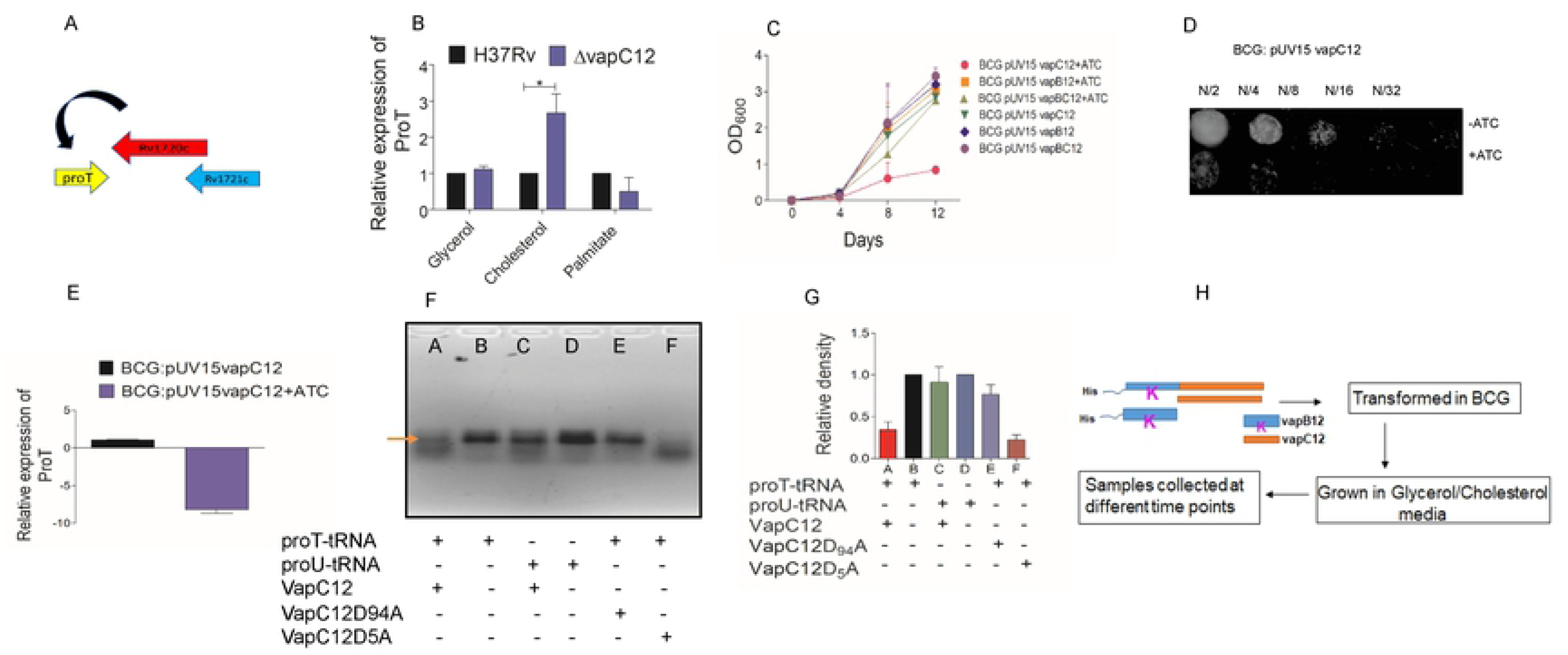
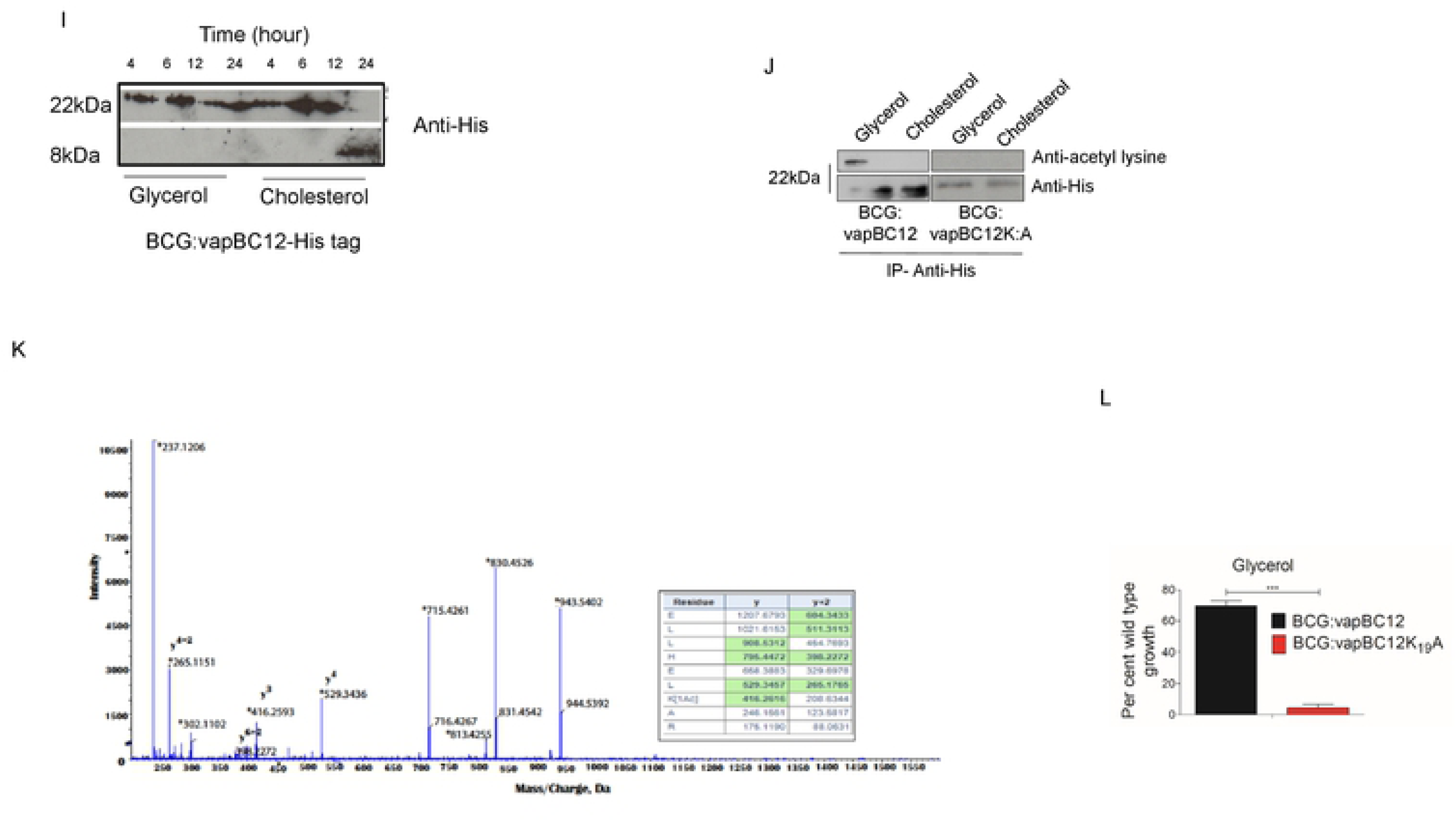
VapC12 ribonuclease toxin targeting proT is essential for cholesterol-mediated growth regulation in Mtb. A) Diagrammatic representation of toxin–antitoxin *vapBC12* locus. B) Relative expression of proT tRNA through qRTPCR in the *vapC12* mutant relative to the wild-type H37Rv strain grown in media containing glycerol, cholesterol, and palmitate as the sole carbon source. C) Growth curve of *M. bovis* BCG strain expressing *vapC12*, *vapB12*, and *vapBC12* in the pUV15-tetO expression system under the tet-inducible promoter in 7H9+OADC media. Anhydrotetracycline (ATc), an inducer of the tet operon, was used at a concentration of 100 ng/mL and replenished every fourth day. D) Two-fold serial dilutions (N/2, N/4, N/8, N/16, N/32) of the log phase growing culture *BCG:pUV15 tetO:vapC12* strain grown in 7H9 broth were spotted on 7H11 agar plates with or without ATc. E) Relative quantification of the transcript levels of proT gene *in BCG:pUV15-tetO:vapC12* grown in 7H9 media with or without ATc by qRT-PCR. F) RNase activity of purified wild-type and mutant VapC12 toxins against in vitro transcribed tRNA substrates. Different wells of the gel denote different combination of tRNA transcript and purified proteins viz; (A) Wild-type VapC12 toxin protein incubated with proT, (B) proT tRNA only with no protein, (C) wild-type VapC12 toxin protein incubated with proU, (D) proU tRNA only, (E) mutant VapC12D_94_A toxin protein incubated with proT, and (F) mutant VapC12D_5_A toxin protein incubated with proT. Each reaction was incubated at 37°C for 3 hours. The products of each of the reaction were run on a 3% agarose gel and visualized by adding ethidium bromide followed by exposure to UV light. G) Relative density of marked RNA bands in Fig 2F quantified using ImageJ. The experiment (2F) was repeated three times, and data plotted represent the mean ± SEM. H) Schematic representation of the protocol for the experiment to demonstrate cholesterol-specific dissociation of the antitoxin. I) Western blot for cholesterol-specific dissociation and degradation of the antitoxin from the toxin–antitoxin complex. The His-tagged antitoxin was tracked using an anti-His antibody in the cell lysate of BCG overexpressing His-tagged antitoxin as a part of the toxin–antitoxin complex. Cell lysates were prepared by sampling cultures grown in both glycerol and cholesterol media at different time points and probed with an anti-His antibody. J) Western blot of the protein lysates prepared from BCG overexpressing toxin-antitoxin locus (VapBC12) with His tagged antitoxin VapB12 and BCG strain with N-terminal His tagged VapBC12 where in lysine residue of AT is converted to alanine (VapB_K:A_C12). Immunoprecipitation was performed using mouse anti-His antibody and probed with rabbit anti-acetyl lysine and anti-His antibodies. To normalize for the amount of the protein, three-fold higher concentration of protein was loaded in the cholesterol-grown BCG sample. K) Mass spectrometry analysis of His-tagged antitoxin protein isolated from BCG overexpressing VapBC12 complex grown in glycerol and cholesterol media. Tryptic digest of immunoprecipitated samples from glycerol and cholesterol grown cultures were analysed by LC-MS/MS (Sciex Triple TOF 5600). Representative MS/MS spectrum of peptide from glycerol grown sample, ELLHELK(Ac)AR was acetylated and displays mass shift corresponding acetylation (m/z 416.26) when compared to the unmodified peptide from cholesterol grown sample. L) Growth curve of BCG overexpressing toxin–antitoxin (*vapBC12*) and lysine mutant (*vapB_K:A_C12*) in a minimal media containing 0.1% glycerol as the carbon source. Bacterial enumeration was performed at day 7 after inoculation by CFU plating on 7H11+OADC plates. Experiment was performed in triplicates, and data plotted represent the mean ± SEM. Data were analysed using unpaired Student’s t test. *P < 0.05, **P < 0.01

**Figure 3:**
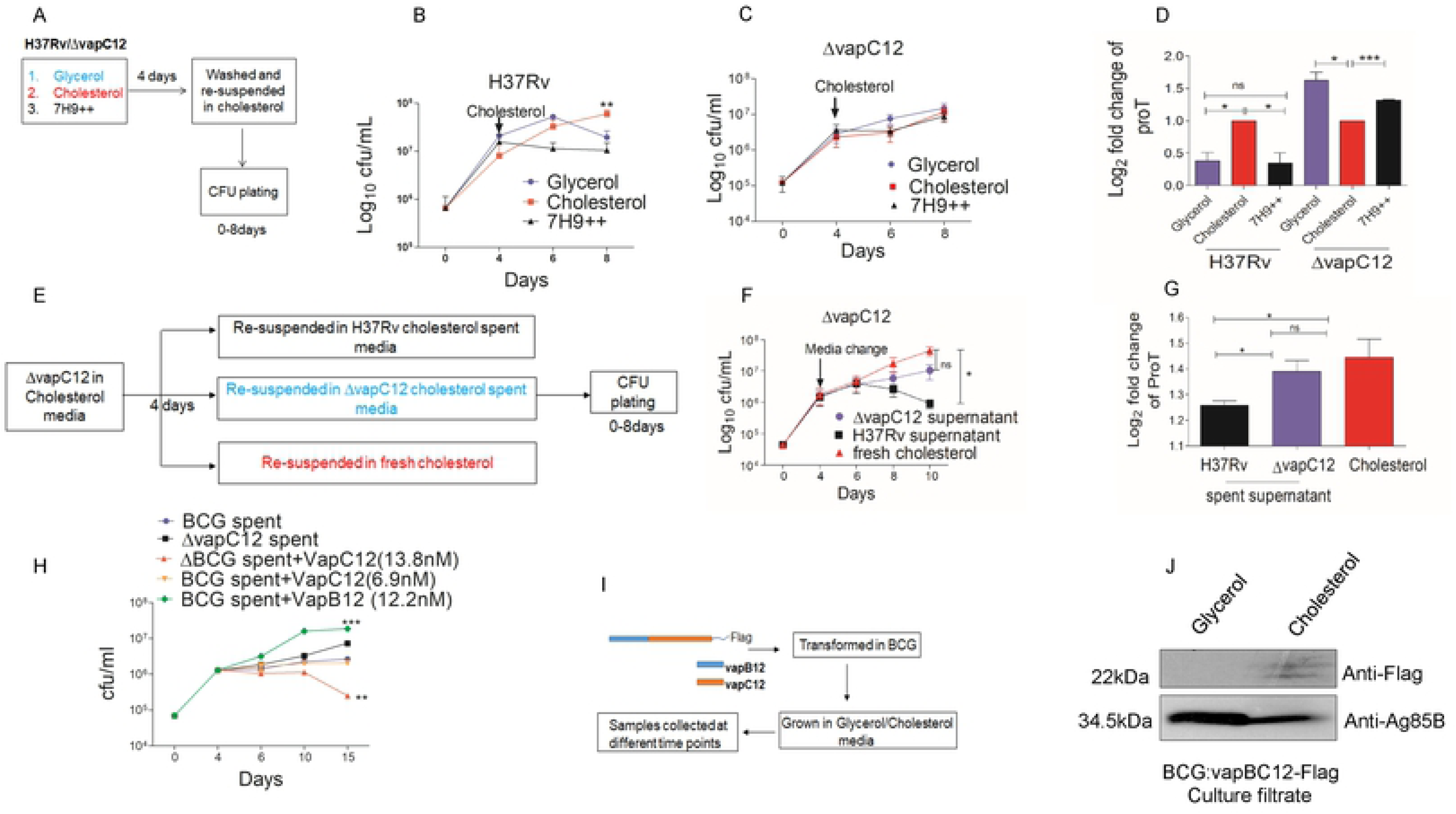
Cholesterol-dependent activation of *vapC12* toxin generates and enriches the persister population in the Mtb culture. A) Schematic representation of the persister enrichment experiment. B) & C) Growth curve of H37Rv and *vapC12* mutant strains grown in 7H9 enriched, 0.1% glycerol and 0.01% cholesterol media for first 4 days and then resuspended in a cholesterol-rich media for subsequent days. Bacterial enumeration was performed by plating cultures on 7H11+OADC plates at various time points. The experiment was performed in triplicates, and data plotted represent the mean ± SEM. Data were analysed using unpaired Student’s t test. *P < 0.05 D) Expression analysis of proT tRNA through qRTPCR in H37Rv and *vapC12* mutant strains at day 8 relative to day 4 of the persister enrichment growth curve (Fig.3B and 3C). E) Schematic representation of growth curves obtained from spent media from wild-type H37Rv and *vapC12* mutant strains grown in cholesterol. F) The *vapC12* mutant strain was grown in a media containing 0.01% cholesterol in triplicate for the first 4 days and then resuspended in spent media from H37Rv, *vapC12* mutant, and fresh cholesterol individually. Bacterial enumeration was performed by plating cultures on 7H11+OADC plates. The experiment was repeated three times, and data plotted represent the mean ± SEM. Data were analysed using unpaired Student’s t test. *P < 0.05 **P < 0.01. G) Expression analysis of proT tRNA through qRTPCR in the *vapC12* mutant strain grown in a cholesterol-spent media at day 10 of the growth curve relative to the culture grown in a fresh cholesterol-rich media at day 4 (Fig. 3F). H) The BCG *vapC12* mutant strain was grown in a media containing 0.01% cholesterol in triplicate for the first 4 days and then resuspended in spent media from wild-type BCG, *vapC12* mutant, and wild-type BCG supplemented with purified VapB12 antitoxin (12.2nm) and VapC12 toxin at two different concentration (6.9nM and 13.8nM). Bacterial enumeration was performed through CFU plating on 7H11+OADC plates. The experiment was repeated three times, and data plotted represent the mean ± SEM. Data were analysed using unpaired Student’s t test. *P < 0.05 **P < 0.01. I) Schematic representation of the experiment to demonstrate that toxin is secreted out in the culture filtrate of BCG. J) Western blot showing the flag-tagged toxin in the culture filtrate of BCG strain overexpressing the toxin–antitoxin complex (VapBC12). The culture filtrate was probed with an anti-flag antibody to detect the toxin protein, anti-Ag85B antibody as a positive control for the secretory protein, and an anti-GroEL1 antibody as a negative control to ensure no lysis of bacterial cells occurred during sample preparation.

**Figure 4:**
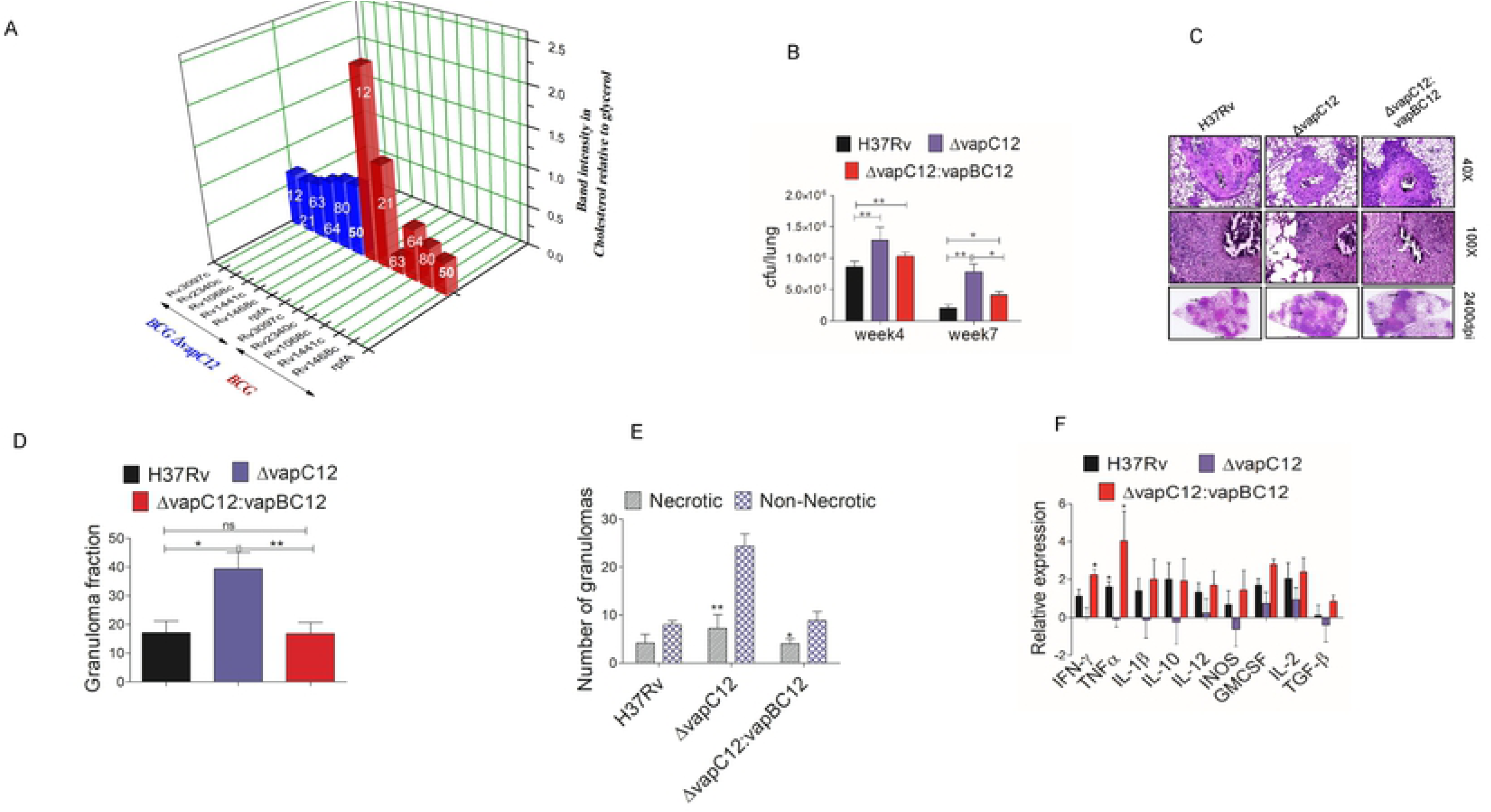
*vapC12*-mediated downregulation of proT-encoded proline-rich proteins are essential for persistence of Mtb in a guinea pig model of infection. A) Relative band intensity representing the expression of His-tagged PE-PGRS and RpfA proteins in BCG and *vapC12* mutant strains grown in glycerol- and cholesterol-rich media. The numbers on each individual bar represent the percentage of ProT codon in that particular protein. The protein lysates were prepared from overexpressed strains with an OD of 0.8–1. The samples were run on SDS PAGE and probed with an anti-His antibody. B) Bacterial load in the lungs of guinea pigs infected with H37Rv, *ΔvapC12*, and *ΔvapC12*:*vapBC12* strains of Mtb. At designated time points, the lungs were homogenized in 4 mL of saline, and ten-fold serial dilutions of homogenates were plated on 7H11+OADC plates. Each group constituted six guinea pigs per time point. Data plotted represent the mean ± SEM. Significant differences observed between groups are indicated. Data were analysed using the Mann–Whitney U test with **P < 0.01 and *P < 0.05. C) Photomicrographs of H&E-stained (40× and 100×) and high-resolution scanning (2,400 dpi) of lung sections from guinea pigs infected with different strains of Mtb at 7 weeks post infection. D) Granuloma fraction of the lung tissue samples of guinea pigs infected with different strains of Mtb, based on the semi-quantitative estimation of the fraction of the lung tissue covered with granuloma. Data were analysed using the Mann–Whitney U test with *P < 0.05 and **P < 0.01. E) Total number of necrotic and non-necrotic granulomas in the lung tissue samples of guinea pigs infected with different strains of Mtb. Data were analysed using the Mann–Whitney U test with *P < 0.05, * P < 0.01, and ***P < 0.001. F) Cytokine profiling of animals infected with H37Rv, *ΔvapC12*, and *ΔvapC12*:*vapBC12* strains of Mtb. RNA was extracted from the spleen of infected animals 7 weeks post infection. The relative expression of cytokines in different groups of animals was quantified through qRTPCR. Data were normalized with the findings of the uninfected group. Data plotted represent the mean ± SEM. Data were analysed using the Mann– Whitney test with *P < 0.05 and **P < 0.01.

### VapC12 ribonuclease toxin targeting proT is essential for cholesterol-mediated growth regulation in Mtb

Because the VapBC family of toxins targets RNAs(*39–41*), the presence of proT-tRNA (proT) gene upstream to *vapC12* gene was intriguing. Thus, we hypothesized that this proT-tRNA can be one of the substrates for the toxin (Fig. 2A). As predicted, we observed a cholesterol specific decrease in the proT transcript levels in wild type strain (Fig. S5) and this cholesterol specific decrease was not observed in a *vapC12* null strain, suggesting, proT tRNA could be one of the major substrates of the VapC12 ribonuclease toxin. In order to further confirm the proT specificity, we quantified the transcript levels of 10 different tRNAs that had a GC-rich anticodon sequences. Surprisingly, we found no cholesterol specific differences in the abundance of any of the tested tRNAs, emphasizing that proT is one of the major substrates of the VapC12 ribonuclease toxin (Fig. S6). To further rule out the role of slow growth rate contributing to the above phenotype, we quantified the proT transcript levels in WT Mtb culture grown in palmitate as a sole carbon source (Fig. S7) and we did not observe any difference in the proT transcript levels (Fig. 2B).

In contrast to the findings of previous studies(*26, 42*), we successfully demonstrated that WT *M. bovis* BCG strain overexpressing the putative toxin gene *vapC12* demonstrated both a decrease in the proT transcript level and a significant growth defect (Fig. 2C, 2D, 2E). Furthermore, the toxin phenotype was reversed if the *vapC12* toxin gene was co-expressed along with its cognate *vapB12* antitoxin gene (Fig. 2C) (*43*). To further validate if proT is the substrate of VapC12 toxin, we generated recombinant His-tagged VapC12 toxin expressed and purified in a heterologous *E. coli* expression system (Fig. S8). When exposed to *in-vitro* transcribed tRNAs, namely proT and proU, the purified recombinant toxin specifically cleaved proT (Fig. 2F, 2G). Furthermore, we mutated two highly conserved aspartate residues D_5_ and D_94_ in the PIN domain of the toxin to alanine (Fig. S9). An aspartate (D) to alanine (A) conversion of the 94^th^ residue of VapC12 toxin failed to cleave the substrate. This inactivation of the toxin by D_94_A substitution may be due to its inability to bind to Mg^2++^, which is a critical cofactor required for its activity(*39, 44*). Although we used proU-tRNA (proU) as our control, we do not rule out the possibility of other tRNAs being a VapC12 substrate. A D_5_A substitution didn’t affect the ribonuclease activity of the toxin.

Activation of the type II TA module is mainly due to the degradation of the corresponding antitoxin. Therefore, to study cholesterol-specific degradation of the antitoxin VapB12, we generated a strain expressing N-terminal His-tagged VapB12 antitoxin and toxin VapC12 at C-terminal. In the presence of cholesterol, the antitoxin (VapB12) protein dissociated from the VapBC12 complex and degrades with time, releasing and activating the cognate toxin (Fig. 2H, 2I). The only lysine residue K_19_ of the antitoxin lost its acetylation in the presence of cholesterol (Fig. 2J) and the de-acetylation of this lysine residue (K_19_) in the antitoxin protein can be the signal for cholesterol-induced degradation of antitoxin and the subsequent activation of VapC12 toxin. This activation modulates cholesterol-specific growth in Mtb.(*45*). We validated this by LC-MS/MS wherein we observed that the lysine residue (K_19_) of the antitoxin protein was acetylated only in protein lysates from glycerol media (Fig. 2K). Surprisingly, the protein coverage of the anti-toxin peptides isolated from Mtb grown in cholesterol was ≥ 95 per cent except for the peptide LHELK with the sequence coverage of ≥50 <95 (Fig. S10). This could be attributed to a cholesterol specific degradation of the anti-toxin (Fig. 2I). To confirm this further, we generated a recombinant *M. bovis* BCG strain overexpressing the antitoxin protein harbouring a lysine to alanine (K_19_A) substitution and as expected due to the absence of the lysine residue the antitoxin could not be acetylated (Fig.2J). This resulted in constitutive degradation of the antitoxin, leading to growth inhibition independent of the carbon source in mycobacteria (Fig. 2L).

### Cholesterol-dependent activation of *vapC12* toxin generates and enriches the persister population in the Mtb culture

In order to further evaluate cholesterol-induced activation of VapC12 toxin and the subsequent slowdown of Mtb growth, a log-phase culture of WT Mtb grown in either an enriched (7H9+OADC) or glycerol media was subsequently exposed to cholesterol, and the effect on bacterial growth (cfu) was assessed (Fig. 3A). Exposure to cholesterol caused a significant reduction in the Mtb growth rate (Fig. 3B), which was dependent on the presence of *vapC12* toxin gene (Fig. 3C). These results indicate a *vapC12*-mediated cholesterol-dependent reduction in Mtb growth. Compared with the glycerol media, the reduction in the growth rate was more prominent in the enriched media. We also found a reduction in the transcript levels of proT-tRNA in cholesterol-exposed Mtb, which was earlier grown in either an enriched media or glycerol media (Fig. 3D), further confirming the finding that the difference in growth is due to toxin-mediated degradation of proline-tRNA. To explore the extracellular role of VapC12 toxin in restricting the growth of fast-growing bacteria in a heterogeneous population (*46, 47*), we suspended a log-phase culture of the Mtb *vapC12* mutant strain separately in the spent media harvested from either the cholesterol-grown WT or *vapC12* mutant strain (Fig. 3E). A decrease in the *vapC12* mutant cfu was observed only in cultures resuspended in the supernatant isolated from the cholesterol-grown WT strain, suggesting that either VapC12 toxin directly or a VapC12-dependent secretory protein selectively enriches the slow growing persister population in Mtb cultures in a cholesterol rich environment (Fig. 3F). The quantification of proT levels in these cultures further suggested that the observed phenotype was indeed due to differences in the toxin-mediated proT cleavage (Fig. 3G). Furthermore, neutralization of the toxin in WT spent media by adding a purified antitoxin resulted in an increase in the mutant CFU, similarly, the addition of purified toxin in spent media from *ΔvapC12* resulted in a dose dependent decrease in CFU of *vapC12* null strain (Fig. 3H). Finally, we demonstrated that VapC12 toxin was only detected in the culture filtrate isolated from cholesterol-grown *M. bovis* BCG overexpressing Flag-tagged VapC12 and not from the glycerol-grown culture (Fig. 3I, 3J).

We next investigated the implications of proT degradation in cholesterol-mediated growth modulation. A Genome-wide *in-silico* analysis was performed to determine the frequency of the pro-tRNA codon in each of the Mtb gene (Fig. S11). Mycobacteria code four designated pro-tRNA (pro-T, pro-Y, pro-U, and pro-X) that incorporate the proline residue to a protein during translation. The results of the in-silico analysis revealed that pro-T and pro-Y encode 85.53% of the total proline incorporated into the Mtb H37Rv proteome, with pro-Y (CCG) and pro-T (CCC) codon usage being 63.6% and 36.4%, respectively (Fig. S11). A list of 136 Mtb genes that had at least 60% of the proline encoded by pro-T tRNA was identified (Table S3). A functional categorization revealed that proline incorporated in the PE-PGRS protein family has a significantly higher proT codon usage. In addition, a gradient in the percentage of proT codon usage was observed (Fig. S12). This led to speculation that the expression of antigenic proteins belonging to the PE-PGRS family, which contain varying number and frequency of proT, are differentially regulated in a cholesterol-rich environment. We hypothesized that through VapC12 toxin-mediated degradation of proT in cholesterol, Mtb downregulates the expression of these antigenic proT-rich PE-PGRS proteins. We selected a set of 5 different PE-PGRS proteins with varying proT codon usage and found that the expression levels of PE-PGRS proteins having a higher frequency of proT decreased in Mtb grown in the cholesterol-rich media compared with that grown in a glycerol-rich media (Fig. 4A, S13). This phenotype was completely dependent on the presence of VapC12 toxin because no media-specific difference was observed in the expression of the aforementioned proteins in the *vapC12*-null strain (Fig. 4A, S13). The factor *rpf A*, one of the five resuscitation-promoting factors (*rpfA-D*) of Mtb, has 53.16% of its proline encoded by proT. These rpf proteins are peptidoglycan glycosidases required for the activation of quiescent bacteria; this is an essential step for the reactivation of TB in a latently infected individual(*48*). In addition, *rpfA* in Mtb has been reported to be secretory through a sec-dependent pathway and speculated to be involved in modulating host during the reactivation process(*49*). Interestingly, we found a *vapC12*-dependent decrease in the expression of *rpfA* in a cholesterol-rich environment. We believe that the cholesterol-dependent regulation of *rpfA* levels by the VapC12 toxin can be an important mechanism through which Mtb sustains latency during infection.

### *vapC12*-mediated downregulation of proT-encoded proline-rich proteins are essential for persistence of Mtb in a guinea pig model of infection

To assess the possible role of *vapC12* in the host, we first infected mouse bone marrow-derived macrophages (BMDM) with WT and *vapC12*-null strains. The null strain demonstrated increased replication in BMDM, which was abolished in the complemented strain (Fig. S13). Of note, the *vapC12*-null strain also showed similar enhanced fitness to replicate under oxidative and nitrosative stress (Fig. S14). Next, we infected guinea pigs with WT, *vapC12*-null, and complemented Mtb strains. At 7 weeks post infection, a higher bacterial load was observed in the lungs of guinea pigs infected with the *vapC12*-null strain compared with animals infected with a WT or complemented strain (Fig. 4B). A similar profile was observed in the spleen (Fig. S16). Histologic examination of the lungs of infected animals at 7 weeks post infection revealed necrotic and non-necrotic lesions with numerous infiltrating macrophages and lymphocytes (Fig. 4C, D). At this time, *vapC12*-null strain infected Guinea pigs displayed an increased number of granulomas (Fig. 4C,4D, S17), which were predominantly of the non-necrotic type (Fig. 4E). This suggests a hypervirulent phenotype of the *vapC12*-null strain. Consistent with this finding, animals infected with the *vapC12*-null strain failed to induce an inflammatory response, as indicated by a decreased mRNA expression of inflammatory cytokines (Fig. 4F). Guinea pigs infected with a complemented strain had a phenotype similar to those infected with a WT strain (Fig. 4C-F).

## Discussion

Chronic infections necessitate the etiologic agent to persist inside the host for extended duration. Mtb remarkably adapts to a very hostile niche by augmenting its ability to thrive inside the host for decades. The pathogen’s ability to modulate host immune response and its capacity to tolerate high concentration of anti-mycobacterial drugs are key to persistence. In order to do so, Mtb senses various stage specific environmental cues and accordingly regulates the expression of various proteins that eventually help the pathogen to attain distinct phenotypes critical for long-term survival. Inside the host, Mtb encounters an extraordinary challenge of surviving on host-derived nutrients and subsequently creating a niche conducive for its own growth. Conversely, the host has developed ways and means to deprive, the unwanted guest, of critical nutrients including the much-needed carbon source. Although, we have earlier demonstrated that the utilization of host cholesterol is essential for disease persistence in tuberculosis, the role of cholesterol utilization and the subsequent mechanism leading to the above phenotype is largely not well defined.

We, in the current study have demonstrated that cholesterol utilization results in an increase in the frequency of generation of antibiotic persisters in mycobacteria. The above phenotype was abrogated in an Mtb ribonuclease toxin *vapC12* null strain. Mechanistically, we also identified Mtb proT tRNA as one of the substrates of the VapC12 ribonuclease toxin and that the toxin mediated modulation of the proT tRNA levels regulate antibiotic persistence in mycobacteria. Finally, using guinea pig model of Mtb infection, we demonstrated that a reduction in the frequency of generation of antibiotic persisters significantly curtailed disease persistence. According to the recently established guidelines on bacterial persistence (*50*), our data suggest that in tuberculosis both antibiotic as well as disease persistence, either individually or in tandem, influence the disease progression and treatment outcomes. Co-evolution for centuries has moulded Mtb to adapt and utilize host-derived fatty acid including cholesterol as a preferred carbon source (*1, 51*). Surprisingly, Mtb does not rely on cholesterol as sole carbon source during infection (*52, 53*), nonetheless, utilization of the host cholesterol has been found to be essential for long-term persistence. Although, nutrient dependent growth modulation is very common (*54, 55*), our data for the first time reported its effect on antibiotic persistence in tuberculosis.

Interestingly, decrease in the expression of esx3 loci, a type VII secretion system critical (*38*)for iron uptake(*36, 56*), during cholesterol utilization suggests that Mtb deprives itself of iron in order to restrict growth under cholesterol rich condition. We also found that the cholesterol exposed Mtb downregulates the expression of genes belonging to the electron transport chain (ETC) resulting in a sharp decline in the intracellular ATP levels. Our data is in line with similar studies implicating lower ATP concentration to disease persistence in several species of bacteria including Mtb (*37, 38, 57, 58*). These findings suggest that the modulation of intracellular ATP levels by a *vapC12* gene-encoded protein might have a role in cholesterol-specific growth modulation in Mtb. Obtaining mechanistic insights into pathways leading to VapC12 toxin-dependent regulation of intracellular ATP levels would be an interesting area for future research. Additionally, an increase in the transcript levels of DosR regulon genes in the cholesterol-rich media suggests that in addition to hypoxia, sensing of intracellular cholesterol by Mtb can possibly trigger the induction of DosR regulon genes in Mtb. Surprisingly, in spite of the cholesterol specific growth differences observed between the WT and the *vapC12* mutant, the differences observed in the transcript levels were very minimal implicating post-transcriptional regulation for the observed phenotype. Our finding demonstrated growth modulation specifically attributed to the abundance of proline-tRNA levels modulated by activation of VapC12 ribonuclease toxin. Studies describing tRNA dependent growth modulation have already been reported (*27, 59, 60*), our study describing mechanism of nutrient dependent regulation of tRNA abundance modulating growth is a novel finding.

Post-translational modifications (PTMs) confer diversity to regulatory mechanisms that controls various cellular pathways. Two of the most extensively studied PTMs are phosphorylation and acetylation. Together, they are known to regulate the stability and activity of proteins in both eukaryotes and prokaryotes. Lysine acetylation is known to regulate various cellular pathways conserved across species including mycobacteria (*45, 61*). Our data also suggest that the cholesterol-mediated growth modulation is triggered by deacetylation of the only lysine residue present in the antitoxin.

Mechanistically, the data suggest that, in cholesterol rich environment, VapC12 toxin enriches slow-growing bacteria by selectively culling fast-growing ones in a heterogeneously growing Mtb culture. These findings unravel a new mechanism through which Mtb regulates the generation and enrichment of persisters when exposed to cholesterol, a carbon source typically available during the persistence stage of tuberculosis infection. Our findings also suggest that the proT-encoded proline-rich proteome of Mtb, including PE-PGRS proteins, have immunomodulatory properties. We predict that these PE-PGRS proteins have relatively higher expression during early stages of infection that blunts the host immune response, resulting in active growth of Mtb inside the host. After the onset of adaptive immunity, increased exposure of Mtb to host cholesterol down regulates the expression of these immunomodulatory proteins, leading to enhanced pro-inflammatory cytokine secretion that triggers granuloma formation and containment of the infection.

Thus, these proT codon rich proteins belonging to the PE-PGRS proteins induce a pro-inflammatory cytokine response and a down-regulation of these proteins during chronic phase of infection is critical for long-term survival and persistence. A recent publication in support of this hypothesis suggest how ubiquitination of one of the proT enriched proline codon protein belonging to the PE-PGRS family was identified as signal for the host to eliminate the pathogen(*62*). Our findings suggest that Mtb through the pathways that we have identified downregulates the expression of such antigens for its long-term survival inside the host. VapC12-mediated proT codon-based differential expression of various Mtb proteins, including PE-PGRS, is a novel finding and can be a pathogen-driven immunomodulatory mechanism critical for the maintenance of the persistence state during mycobacterial infection. While, pathogen rewiring their metabolic pathways for disease persistence is quite well studied (*63, 64*), there are limited studies pertaining to the modulation of host immune response by temporospatial expression of Mtb surface antigens contributing towards disease persistence. Our study suggests that both the growth modulation and differential expression of surface antigens by Mtb together contribute towards disease persistence. This information could be used for designing better and more efficient vaccine against tuberculosis.

In light of our current findings, it will be very intriguing to study the role of PE-PGRS proteins in modulating the host response and their role in the disease progression during Mtb infection. Furthermore, functional characterization of proT tRNA-encoded proline-rich proteins and their implications in the stage-specific replication and growth rate of Mtb inside the host should be explored.

The findings support our hypothesis that the VapC12 toxin acts as a molecular switch that regulates growth in the presence of cholesterol. Because an actively growing Mtb culture is always heterogeneous and has individual bacteria growing at different rates, the rate of growth is directly proportional to the level of the VapBC12 TA protein accumulated in the cytoplasm. The fate of each bacterium after cholesterol exposure is dictated by the intracellular concentration of the activated toxin generated in an individual cell. Depending on the toxin level, bacteria either gets eliminated or slows down, resulting in an enrichment of the persister population when exposed to a cholesterol-rich environment (Fig. S18). Furthermore, the extracellular presence of this toxin ensures clearance of any rapidly dividing mutant bacteria generated due to spontaneous incorporation of a genetic lesion. This is the first study to identify a novel mechanism of cholesterol-dependent stochastic enrichment of slow-growing Mtb during mycobacterial infection.

These findings will help identify novel mechanism of generation of antibiotic persistence and define targets against persister population. Approaches targeting persister population will enhance the rate of clearance of the pathogen resulting in a significant reduction in the duration of treatment. This will help in significantly reducing the risk associated with the current extended regimen extending from six months to two years. So, we have empirically demonstrated that both antibiotic and disease persistence contributes towards chronic Mtb infection and targeting pathways essential for both could potentially shorten the treatment regimen. The current finding holds significance as better understanding of the disease persistence and targeting Mtb persister population as a therapeutic strategy will open new paradigms in tuberculosis treatment.

## Acknowledgement

This study was supported by the India-Singapore grant jointly sponsored by the Department of Science and Technology (DST), India and A*STAR, Singapore to AKP (#INT/Sin/P-08/2015) and AS (#1518224018), respectively. Ramalingaswami fellowship (#BT/RLF/Re-entry/45/2010) and RGYI grant (#BT/PR6556/GBD/27/459/2012) from Department of Biotechnology, Govt. of India and Intramural funding by THSTI to A.K.P is acknowledged. S.T. is funded by research fellowship from ICMR (2012/HRD-119-31498). We thank the Director, ICGEB, and staff TACF, ICGEB, New Delhi for use of the BSL3 facility. Assistance of Mr. Sudesh Rathaor for animal experiments, Mr. Surjeet Yadav for lab maintenance and Dr. Ashok Mukherjee for histopathological analysis is acknowledged.

## Authors Contributions

A.K.P and S.T. designed the experiments. S.T and M.P. performed the experiments. C.S. purified the recombinant proteins. R.K. and D.D. performed bioinformatic analysis. J.L, D.C, M.P, A.S performed RNA sequencing and analysed the data. R.G. performed and analysed Mass spectrometry. S.T, A.S and A.K.P analysed the data. S.T. and A.K.P wrote the manuscript with contributions from all co-authors. A.K.P. conceived the idea and supervised the overall study.

## Materials and Methods

### Bacterial Strains and culture

*Mycobacterium tuberculosis* mutants were derived from strain H37Rv using homologous recombination between the suicide plasmid and bacterial genome. 1000bp flanking regions of the target gene, Rv1720c (*vapC12*) were cloned in pJM1 suicide vector and electroporated in H37Rv competent cells using standard protocol from *Mycobacterium tuberculosis* protocols (Tanya Parish & Neil G. Stroker). The strains were maintained on Middlebrook 7H11 agar or 7H9 broth (Difco^TM^ Middlebrook 7H11 Agar,283810 and 7H9 broth,271310) supplemented with 10% OADC enrichment. Hygromycin were added at 50ug/ml respectively. To complement *vapC12* mutant, the lox-flanked chromosomal hygromycin-resistance gene was excised by expression of Cre recombinase. This strain was transformed with pJEB402 harboring the Rv1720c-1721c (*vapBC12*) genes. For growth on defined carbon sources, strains were grown in “minimal media” (0.5 g/L asparagine, 1 g/L KH_2_PO_4_, 2.5 g/L Na_2_HPO_4_, 50mg/L ferric ammonium citrate, 0.5g/L MgSO_4_*7H2O, 0.5mg/L CaCl_2_, 0.1 mg/L ZnSO_4_) containing 0.1% glycerol (v/v) or 0.01% cholesterol (w/v) and 50mg/ml of Sodium palmitate. Growth was determined by CFU plating at different time points on 7H11 with 10 percent OADC plates.

### Growth curve

The log phase cultures of wild type H37Rv, *ΔvapC12*, and *ΔvapC12:vapBC12* strains were washed with PBST twice and inoculated in minimal media with 0.1 per cent glycerol and 0.01 per cent cholesterol respectively at an absorbance of 0.005. The aliquots of the cultures were taken at different time points and plated on 7H11+OADC plates for bacterial enumeration.

### Resazurin based metabolic activity assay

The log phase culture of wild type H37Rv, *ΔvapC12* and *ΔvapC12:vapBC12* strains of 0.5 OD were washed with PBST twice, and OD_600_ was set to 0.05 in glycerol and cholesterol media. These cultures were serially diluted in the respective media in 96 well plate. The experiment was done in duplicate and both the plates were incubated at 37^0^C for five days before PrestoBlue cell viability reagent (Invitrogen catalogue no. A13261) was added to each well in one set of the plates. The plates were incubated for another two days. The fluorescence read-out of plate with PrestoBlue was taken at 570/585nm using a Synergy HTX Multi-Mode Microplate Reader. For bacterial enumeration in each well CFU plating was done from plate with no Prestoblue. For determining the average metabolic activity, the total fluorescence recorded was normalized for the number of bacteria in the corresponding well.

### Antibiotic kill curve

The log phase culture of wild type H37Rv, *vapC12* mutant and *ΔvapC12:vapBC12* strains grown in 7H9 enriched media were washed with PBST twice and inoculated in glycerol and cholesterol media at an absorbance of 0.05. The cultures were allowed to grow for 4 days before being treated with 5X MIC of rifamycin. Bacterial enumeration was performed through CFU plating of cultures on 7H11+OADC plates at various time points. The kill curve was plotted by calculating the percent survival.

### In-vitro stress assay

The log phase culture of wild type H37Rv and *vapC12* mutant strains were washed with PBST twice and inoculated at an absorbance of 0.1 in 7H9 enriched media for each stress condition keeping an un-treated control. The survival was plotted by CFU plating at different time points post treatment for different stress conditions viz., Oxidative (5mM H_2_O_2_ for 6hrs), Nitrosative (200μM DETA-NO for 24hrs).

### Bone marrow derived macrophages

Bone marrow-derived macrophages (BMM) were isolated by culturing bone marrow cells from C57BL6 mice in DMEM containing 10% FBS, 2mM glutamine, 10% L929-conditioned media, and 10µg/ml ciprofloxacin for 5 days. Approximately 24 hours prior to infection, differentiated BMM were detached and seeded on a 24-well tissue culture plate at 5x 10^5^ cells/well in the same media lacking antibiotic. Macrophages were infected with different strains of *M. tuberculosis* at a MOI of 1 for 4hrs at 37°C and 5 percent carbon dioxide. Extracellular bacteria were removed by washing three times with warm PBS. Intracellular bacteria were quantified by lysing the cells with 0.01% Triton-X100 (Sigma, CAS:9002-93-1) at the indicated time points and plating dilutions on 7H11 agar.

### RNA sequencing material and methods

Log phase cultures of H37Rv and *ΔvapC12* were washed with PBST twice and inoculated in Glycerol and Cholesterol media at an absorbance of 0.005. RNA was isolated from the cultures at day4 using Qiagen RNaeasy Minikit according to manufacturer’s protocols (Qiagen 74104). The RNA was DNase treated using Turbo DNA free kit using manufacturer’s protocol (Thermo Fischer scientific) to remove any genomic DNA contamination. All Mycobacterial total RNAs were analyzed using an Agilent Bioanalyser (Agilent, Santa Clara, CA, USA) for quality assessment with RNA Integrity Number (RIN) range of 5.6 to 9.7 and a median of 7.5. Ribosomal RNA (rRNA) were depleted from 500ng of bacterial RNA using RiboMinus^™^ Bacteria transcriptome isolation kit (Invitrogen Thermo Fisher Scientific Waltham, MA, USA), according to manufacturer’s protocol. cDNA libraries were prepared from the resultant rRNA depleted RNA and 1 ul of a 1:500 dilution of ERCC RNA Spike in Controls (Ambion® Thermo Fisher Scientific, Waltham, MA, USA) using Lexogen SENSE Total RNA-Seq Library Prep Kit (Lexogen GmnH, Vienna, Austria) according to manufacturer’s protocol except with 21 PCR cycles. The length distribution of the cDNA libraries was monitored using a DNA High Sensitivity Reagent Kit on the Perkin Elmer Labchip (Perkin Elmer, Waltham, MA, USA). All samples were subjected to an indexed paired-end sequencing run of 2×51 cycles on an Illumina HiSeq 2000 system (Illumina, San Diego, CA, USA) (16 samples/lane). Raw reads (FASTQ files) were mapped to the *M. tuberculosis* H37Rv (GenBank accession AL123456) using bowtie2 using default parameters. The gene counts were then counted using featureCounts (part of the Subread package) using the genome annotations provided in the GenBank file. The gene counts were then used in DESeq2 for differential gene expression analysis. Multiple testing correction was performed using the method of Benjamini and Hochberg. P values < 0.05 were deemed to be statistically significant. Computations were done using the R statistical language version 3.3.1.

### Quantitative RT-PCR

Comparative qRT-PCR was done from RNA isolated from various strains in different experimental conditions. The RNA isolation was done from culture using RNaeasy Minikit according to manufacturer’s protocols. (Qiagen 74104). The RNA was DNase treated using Turbo DNA free kit according to manufacturer’s protocol (Thermo Fischer scientific) for making cDNA using Accuscript hi-fidelity cDNA synthesis kit (Agilent). The qRT-PCR was set up using brilliant III ultra-fast SYBRgreen qPCR master mix in Mx3005P qPCR system Agilent. The primers used are listed in Table S4. The data analysis was done using MxPro software.

### VapC12 expression and protein purification

*vapC12* was cloned in pET28a vector using primers sequence listed in Table S4. The SDM mutants D5:A and D94:A were generated by dpn1 treatment. The clones were transformed in *E. coli Rosetta* cells. Briefly, the overnight culture was inoculated in 1 L of fresh LB media (1:100) supplemented with 100 µg/mL of kanamycin and allowed to grow until the optical density at 600 nm reached ∼0.5. The culture was then induced with 1 mM IPTG and allowed to grow for overnight at 37°C. Cells were harvested by centrifugation at 6000 × g for 10 minutes and checked for expression of wild-type or mutant rRv1720c by SDS–PAGE. Most of the target protein was present in the pellet as inclusion bodies (IBs). Isolation of pure IBs containing rRv1720c was performed by sonication and several washing steps (Singh et al., 2005).

Purified rRv1720c IBs (1 mL) were solubilized in 9 mL buffer [[50 mM Tris–HCl, pH 8.0, 300 mM NaCl, 10 mM β-mercaptoethanol, 8 M Urea] and incubated at room temperature for 1 h, followed by centrifugation at 15,000 × g for 20 m at 10°C. The supernatant obtained post-centrifugation was used for purification recombinant Rv1720c protein, by immobilized metal ion affinity chromatography (IMAC) using HisTrap FF column (GE Healthcare Buckinghamshire, UK), in denaturing condition. Protein was eluted using buffer [50 mM Tris–HCl, pH 8.0, 300 mM NaCl, 10 mM β-mercaptoethanol, 8 M Urea, 250 mM Imidazole]. Denatured purified protein was refolded by diluting it in a pulsatile manner in refolding buffer [100 mM Phosphate buffer, pH 6.4, 300 mM NaCl, 5 mM β-mercaptoethanol] at 4°C with constant stirring. The refolded target protein sample was centrifuged at 24,000 × g for 30 m at 4°C and the supernatant containing refolded active protein was concentrated and dialyzed three times against buffer [100 mM Phosphate buffer, pH 6.4, 300 mM NaCl, 5 mM β-mercaptoethanol]. The final buffer exchange of protein to buffer [100 mM Phosphate buffer, pH 6.4, 300 mM NaCl, 10% glycerol] was performed by PD10 desalting column (GE Healthcare Buckinghamshire, UK), as per manufacturer’s protocol. Protein was quantitated by bicinchoninic acid assay (Thermo Scientific Pierce, Rockford, IL, USA) and analyzed with SDS-PAGE, and confirmed by Western blotting, using anti-His monoclonal antibody (Cell Signaling Technology, Inc., MA, USA). Similar protocol was followed for purification rRv1720cD_5_A and rRv1720cD_94_A.

Also, *vapB12* cloned in pET28a *E. coli Rosetta* cells, was purified from supernatant of lysed induced cells using Hispur Cobalt purification kit, 3ml (Thermo Scientific, 90092).

### In-vitro transcription of tRNA

The tRNAs were transcribed using Megascript kit (Invitrogen, AM1334) according to manufacturer’s protocol, which is followed by phenol chloroform extraction and isopropanol precipitation for purified transcript. The primers used for in-vitro transcription are listed in Table S4.

### In-vitro tRNA cleavage assay

In-vitro RNA cleavage assay was performed using tRNAs produced via T7 transcription. Cleavage reactions using 3 pmol of tRNA were incubated with 30 pmol of recombinant proteins (rRv1720c, rRv1720cD_5_A and rRv1720cD_94_A) and at 37°C for 3hours in tRNA cleavage buffer (10 mM HEPES (pH 7.5), 15 mM potassium chloride, 3 mM magnesium chloride and 10% glycerol).The samples were mixed with 2X formamide gel loading buffer (95% w/v formamide, 50 mM EDTA) and incubated at 95°C for 5 mins before running on 3% agarose gel. The bands were visualized using ethidium bromide and exposing the gel to UV light.

### Persister enrichment assay

The log phase culture of wild type H37Rv and *ΔvapC12* strains were washed with PBST and an OD of 0.005 was set in 7H9 enriched media, Glycerol and cholesterol media. The cultures were washed after four days and fresh cholesterol media was added in all the tubes. Bacterial enumeration was done by CFU plating on 7H11+OADC plates at different time points.

### Spent media preparation

The log phase culture of H37Rv and *ΔvapC12* were washed with PBST and inoculated at an OD of 0.005 in cholesterol media, the cultures were grown for a week at 37°C in an incubator shaker. The cultures were pelleted down and the supernatant was filtered through 0.2μm filter. The spent cholesterol supernatant of both the strains were used for further experiments.

For preparing heat inactivated supernatant, the spent media was exposed to 95^0^C for 30 minutes before being added to the cultures.

### Spent media experiment

The log phase culture of *ΔvapC12* was washed and inoculated in cholesterol media at an OD of 0.005 in triplicates, the cultures were grown for 4 days after which the culture were washed with PBST and re-suspended in spent H37Rv cholesterol supernatant, H37Rv cholesterol spent supernatant media supplemented with purified toxin or antitoxin, *ΔvapC12* cholesterol spent supernatant according to the experiment, and fresh cholesterol. Bacterial enumeration was done by CFU plating on 7H11+OADC plates at different time points.

### Culture filtrate experiment

*vapB12*(Anti-toxin) and *vapC12*(Toxin) were cloned as an operon in pMV261 kanamycin vector with C-terminal flag tag using primers listed in Table S4. The clone was electroporated in *M.bovis* BCG and maintained in 7H9 enriched media. The log phase culture of C-terminal flag tagged BCG*:vapBC12* was washed with PBST twice and inoculated in glycerol and cholesterol media at an OD of 0.5 and allowed to grow for 48 hours. Cultures were pelleted down and supernatant was filtered through 0.2μm filter. The supernatant was precipitated with 5 percent TCA overnight at 4°C and centrifuged in oak ridge tubes at 12000rpm for 20 minutes at 4^0^C. The pellet thus obtained was washed twice with ice-cold acetone and was allowed to dry and re-suspended in 1X laemmli buffer. The samples were run on 15 percent PAGE and developed with rabbit anti-flag antibody (Sigma, catalogue no. F7425). The samples were also blotted against Ag85B antibody (Abcam, ab43019) and Hsp65 antibody (A kind gift from Dr. Vinay K. Nandicoori) as a positive and negative control respectively.

### Anti-toxin degradation experiment

*vapBC12* operon with a N-terminal His tag was cloned in pMV261 vector. The clone was electroporated in *M. bovis* BCG. The culture was maintained in 7H9 enriched media. The log phase culture was washed with PBST twice and inoculated in glycerol and cholesterol media at an OD of 0.5. Aliquots of culture were taken out, washed and lysed in PBS at different time points. (4hours, 6hours, 24hours,48hours) post inoculation. The lysates were run on a 15% PAGE and developed using monoclonal anti-His antibody (Biospecs, BTL1010) with super signal west femto maximum sensitivity substrate (Thermo Scientific, 34095).

### PE-PGRS/rpf A blots

All the selected PE-PGRS genes along with the *rpfA* (with high proT and high proY codon usage) were cloned in pMV261 using primers listed in Table S4. The proteins were translationally fused with His-tag at the N-terminal by incorporating bases encoding His-tag in Forward cloning primers. The clones were transformed in BCG and the cultures were maintained in 7H9 enriched media. The log phase culture of the constructs was washed twice with PBST and inoculated at an OD of 0.005 in glycerol and cholesterol media. The cultures were allowed to grow till an OD of 0.8 and then were pelleted, washed and lysed in PBS. The samples were run on a 10 % SDS-PAGE gel and developed with anti-His antibody (Biospecs, BTL1010).

### In-vivo animal experiments

The animal experiments were approved by the animal ethics committee of ICGEB (approval no. IAEC/THSTI/2015-1)The animal experiments were performed in accordance with guidelines of Committee for purpose of control and supervision of experiments on animals (CPCSEA, Govt. of India). The pathogenicity of *vapC12* mutant strain was checked by infecting 3-4 weeks old Hartley strain of female guinea pig (200-300gm). The guinea pigs were infected with 100 bacilli of each strain via aerosol route using log phase culture (OD 0.8 to1) of various Mtb strains. For CFU analysis animals were sacrificed and tissues were homogenized at day1, week 4 and week 7 post infection, plating was done on 7H11+OADC plates. For histopathological analysis, lung sections were fixed in 10 percent formalin and stained with hematoxylin and eosin. The tissue samples were coded and evaluated for granulomatous organization by a pathologist who has no prior knowledge of the samples. All granulomas in each section were scored and the scores were added up to obtain a total granuloma score of lungs of each animal. In addition, sections were semi-quantitatively assessed for percentage of the section occupied by granuloma and this was expressed as granuloma fraction.

### Cytokine profiling and RNA isolation from spleen

Single cell suspension of splenocytes was made by passing the spleen through a cell strainer (0.45µ). The back of the syringe plunger was used to macerate the cells through the filter. The cell pellet was incubated with RBC lysis buffer for 4-5 mins after which the pellet was used for extracting RNA. RNA isolation was done from spleen of un-infected and guinea pigs infected with various strains Mtb at week 7 post infection using Rnaeasy Minikit according to manufacturer’s protocol. cDNA synthesis and qRT-PCR were done as described previously using primers listed in Table S4.

### Lysine SDM generation

The SDM mutant of *vapB12* wherein (lysine)K_19_ to (Alanine)A_19_ was mutated to was generated by Dpn1 enzyme treatment. The *vapBC12*His pMV261kan plasmid was used as a template for PCR amplification using SDM primers listed in Table S4. The PCR product was PCR purified and treated with Dpn1 enzyme for 4 hours at 37° C. No template and no Dpn1 treatment controls were also taken. All the reactions were transformed in *E.coli* XL-1 blue competent cells. The construct was sequenced and electroporated in *M. bovis* BCG competent cells.

### Immunoprecipitation

The BCG:*vapBC12* His tagged pMV261 and BCG:*vapB_K19A_C12* His tagged cultures were maintained and grown in 7H9 enriched media, the log phase cultures of the strains were washed twice with PBST and resuspended in glycerol and cholesterol media at an OD of 0.1. After 48 hours the cultures were pelleted and lysed in 1XPBS. The protein quantification of the lysates was done using Pierce BCA Protein Assay Kit - Thermo Fisher Scientific according to manufacturer’s protocol. 1mg of the lysate was incubated with 1:200 dilution of mouse anti-His antibody (Biospecs BTL1010) and incubated overnight at 4° C on a rocker. The Ag-Ab complex was incubated with protein G agarose beads and kept at 4 ° C on a rocker for 8-10 hours. After the incubation the supernatant was collected and beads were washed with 1X TBST thrice. The final elution was done with 100mM glycine pH2.0, which was neutralized later with tris pH 8.8. The eluted samples were run on the gel and blotted with rabbit anti-His (Santa Cruz, H-15:sc-803) and rabbit anti acetylated lysine antibody (Cell signaling, 9441L).

### Sample processing protocol for mass spectrometry

BCG strain (*vapB_K:A_C12*), overexpressing the toxin–antitoxin complex where antitoxin is His tagged with site-directed mutagenesis (SDM) converting lysine to alanine (K_19_to A_19_). Log phase culture of BCG strain (*vapB_K:A_C12*) was washed with PBS and inoculated in minimal media with 0.1 percent glycerol and minimal media with 0.01 percent cholesterol. The cultures were allowed to grow for 48 hours and cell lysate was prepared. The Immunoprecipitation was performed using an anti-his antibody (BTL1010).

In-solution digestion was carried out for 10ug of proteins form each condition. The samples were subjected to reduction and alkylation using 5mM dithiothreitol (DTT) (60C for 45 min) and alkylation using 10 mM iodoacetamide (IAA). Trypsin (Gold mass-spectrometry trypsin; Promega, Madison,WI) digestion was carried out at 37C for 10-12 h. The peptides were vacuum-dried and stored at - 80C until LC-MS/MS analysis.

### LC-MS/MS analysis

All fractions were evaluated by 5600 Triple-TOF mass spectrometer which is directly linked to reverse-phase high-pressure liquid chromatography Ekspert-nanoLC 415 system (Eksigent; Dublin, CA). 0.1% formic acid in water was used as mobile phase A and mobile phase B is 0.1% formic acid in ACN. All fractions were eluted from the analytical column at a flow rate of 250 nL/min using an initial gradient elution of 10% B from 0 to 5 min, transitioned to 40% over 120 min, ramping up to 90% B for 5 min, holding 90% B for 10 min, followed by re-equilibration of 5% B at 10 min with a total run time of 150 min. Peptides were injected into the mass spectrometer using 10 μm SilicaTip electrospray PicoTip emitter. Mass spectra (MS) and tandem mass spectra (MS/MS) were recorded in positive-ion and high-sensitivity mode with a resolution of ∼35,000 full-width half-maximum. Before running samples to mass spectrometer, calibration of spectra occurred after acquisition of every sample using dynamic LC–MS and MS/MS acquisitions of 100 fmol β-galactosidase. The ion accumulation time was set to 250 ms (MS) and to 70 ms (MS/MS). The collected raw files spectra were stored in .wiff format.

### Mass spectrometry data analysis

All raw mass spectrometry files were searched in Protein Pilot software v. 5.0.1 (SCIEX) with the Paragon algorithm. For Paragon searches, the following settings were used: Sample type: Identification; Cysteine Alkylation: Iodoacetamide, Digestion: Trypsin; Instrument: TripleTOF5600. Species: H37Rv maximum allowed missed cleavages 1, Search effort: Thorough ID; Results Quality: 0.05. Only peptides with a confidence score of > 0.05 were considered for further analysis and bias correction was automatically applied. False discovery rate analysis was also performed through decoy database. Carbamidomethylation (C) was used as a fixed modification. The peptide and product ion tolerance of 0.05 Da was used for searches. The output of this search is a. group file and this file contains the following information that is required for targeted data extraction: protein name and accession, cleaved peptide sequence, modified peptide sequence, relative intensity, precursor charge, unused Protscore, confidence, and decoy result.

### Mass spectrometry data submission

The mass spectrometry data obtained from this study has been submitted to public data repositories. The raw proteomics data has been deposited to the ProteomeXchange Consortium via the PRIDE partner repository with the dataset identifier PXD014323. The data also submitted to Massive database at the following link- http://massive.ucsd.edu/ProteoSAFe/status.jsp?task=f56cfab246304977aa56f3e54aa9193

### Codon usage

The bioinformatic analysis was done for determining the codon usage of proT and proY in each gene belonging to all ten functional categories. All the data and the code used for the analysis is given the link below. https://github.com/ddlab-igib/mtb-codon-usage

### ATP estimation

Log phase culture *M. bovis* BCG wild type and *ΔBCGvapC12* strains were washed with PBST twice and inoculated in 0.1 per cent glycerol and 0.01 per cent cholesterol media at an absorbance of 0.005. The aliquots of the cultures were taken at day 5 for ATP estimation. 1ml of each of the culture was pelleted down and resuspended in 0.5ml of PBS followed by heat lysis of cultures at 98°C for ten minutes. ATP was estimated from bacterial lysates using Bac Titer-Glo^TM^ Assay kit from Promega using manufacturer’s protocol. The protein estimation was done in the lysate using Pierce BCA Protein Assay Kit - Thermo Fisher Scientific according to manufacturer’s protocol.

### Statistical analysis

Statistical analysis and graph generation was done using Prism 5 software (Version 5.01; GraphPad Software Inc., CA, USA). For normally distributed data, un-paired student t-test was performed on the means of at least three independent experiments. For animal experiment data analysis Mann Whitney test was performed. P values of less than 0.05, 0.01 and 0.005 were represented to be significantly different as *, ** and *** respectively.

## Supplementary legends

**S1.**
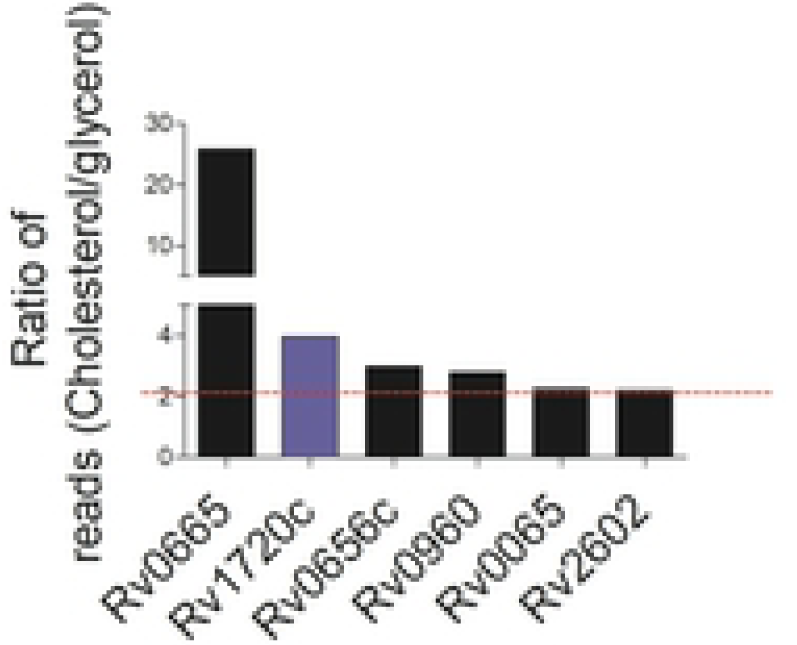
TraCS data representing transposon mutants of *vapC* genes that were overrepresented by more than 2-fold in a cholesterol-rich media compared with a glycerol-rich media, as calculated by number of reads detected per TA insertion site(*32*).

**S2.**
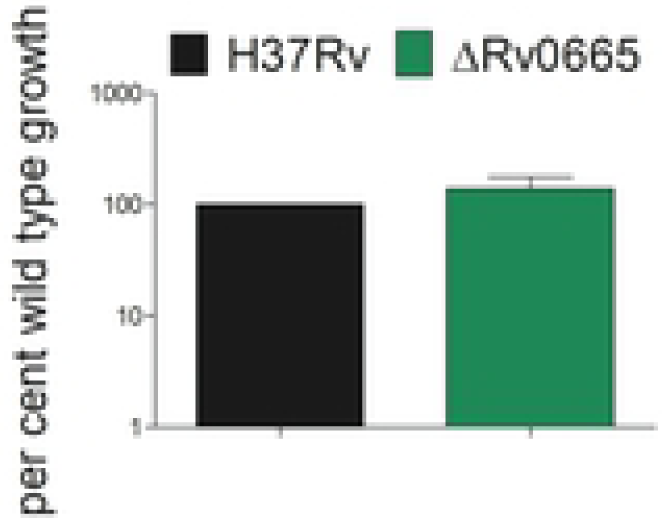
Log-phase cultures of H37Rv and ΔRv0665 (*ΔvapC8*) grown in the 7H9 enriched media were washed with PBS+tyloxapol and resuspended in a media containing 0.01% cholesterol at an absorbance of 0.005. Percent survival of *ΔvapC8* relative to wild-type H37Rv was estimated by plating the culture at day zero and day 8 post inoculation.

**S3.**
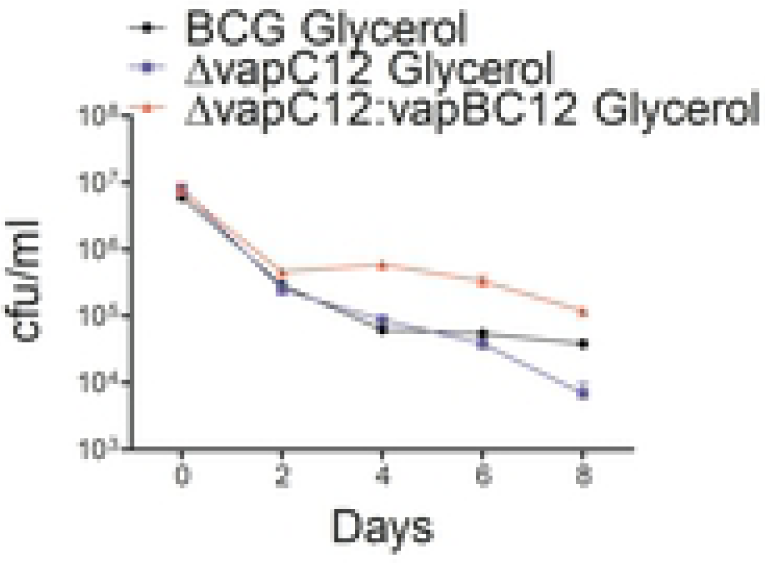
The kill curve of *M. bovis* BCG, BCGΔ*vapC12* and BCGΔ*vapC12:vapBC12* grown in the glycerol-rich media. Log-phase cultures of strains were washed with PBS-tyloxapol and inoculated at an absorbance of 0.05. The cultures were allowed to grow for 4 days before being treated with 5× MIC of rifamycin. Bacterial enumeration was performed by plating cultures on 7H11+OADC plates at various time points. The kill curve was plotted by plotting CFU. The experiment was repeated three times, and data plotted represent the mean ± SEM. Data were analysed using unpaired Student’s t test.

**S4.**
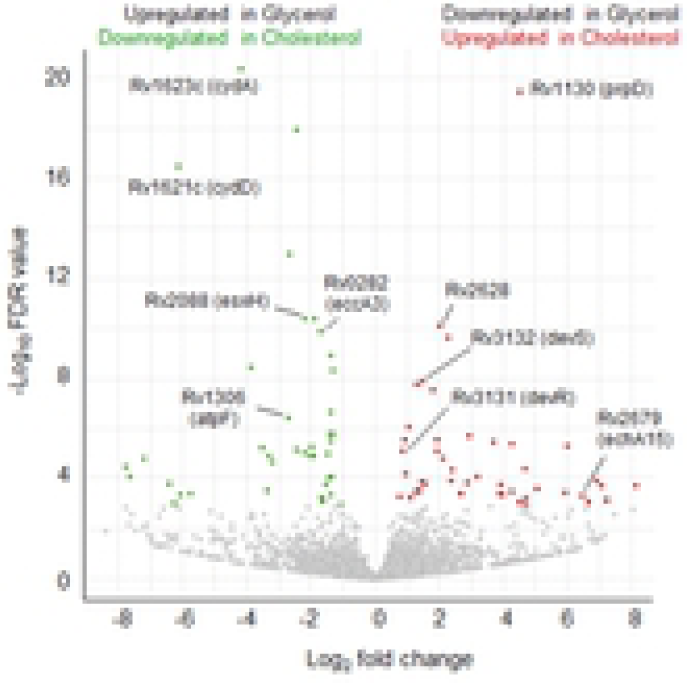
Volcano plot of differentially expressed genes in H37Rv grown in the cholesterol-rich media relative to the glycerol-rich media. Transcriptome of Mtb exhibited 39 downregulated and 45 upregulated genes in the cholesterol-rich media relative to the glycerol-rich media.

**S5.**
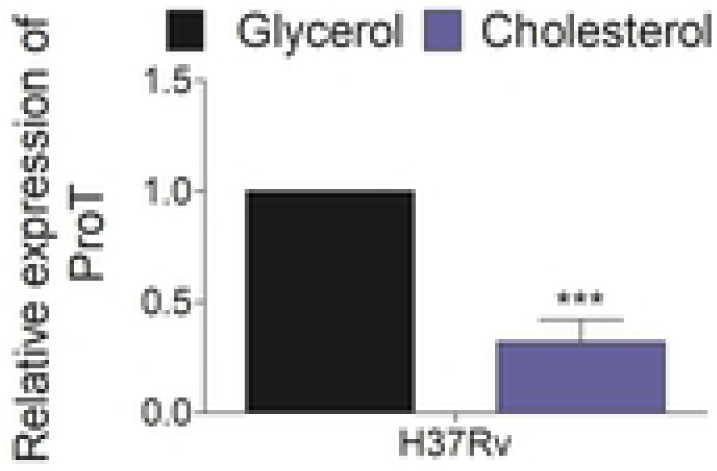
Relative expression of proT tRNA through qRTPCR in wild-type H37Rv strain grown in media containing glycerol and cholesterol as the sole carbon source.

**S6.**
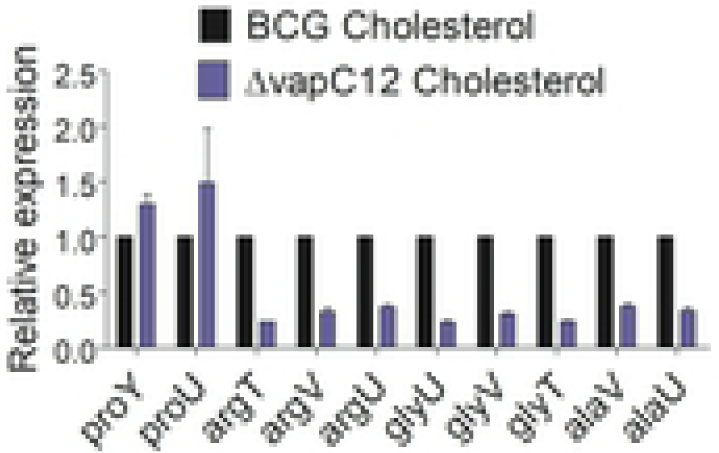
Relative expression of 10 tRNAs through qRTPCR in wild-type H37Rv and *ΔvapC12* grown in the cholesterol-rich media.

**S7.**
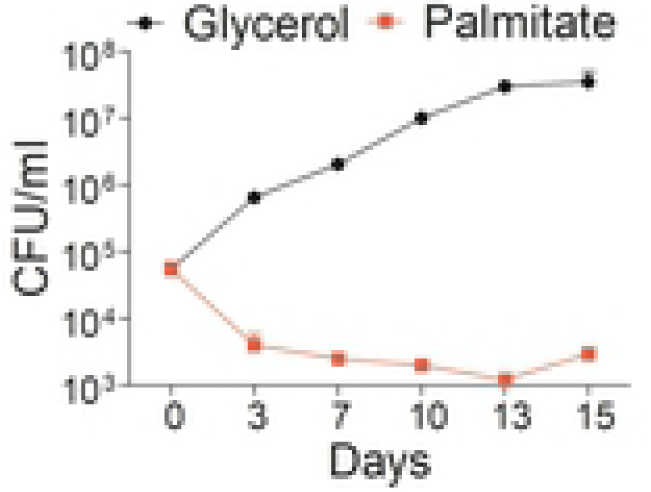
The growth curve analysis of BCG grown in a minimal media supplemented with 0.1% glycerol and 50mg/ml palmitate. Log-phase cultures of wild type BCG grown in 7H9 media enriched with OADC was washed with PBS-tyloxapol and resuspended in respective media at an absorbance of 0.005. Growth was estimated by CFU plating on 7H11+OADC plates at different time points post inoculation, and colonies were counted after 3 weeks of incubation of plates at 37°C. Experiments were performed in triplicates, and data represent the mean ± SEM.

**S8.**
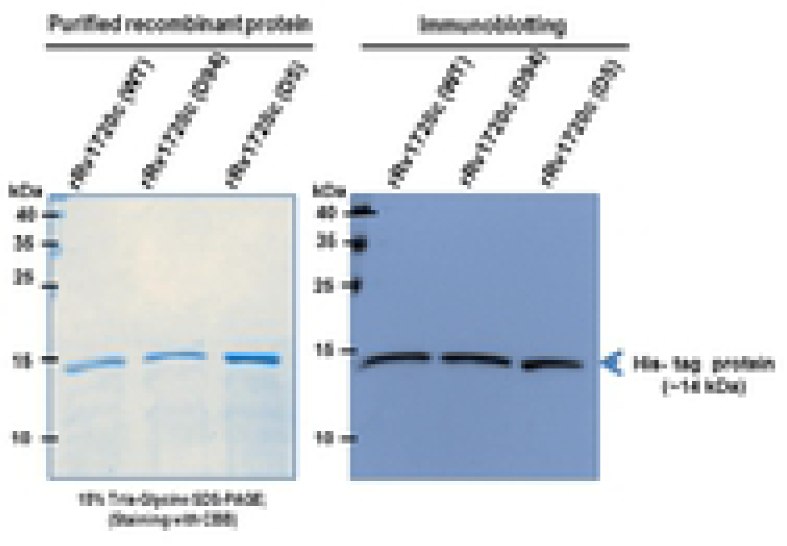
Purified recombinant VapC12, VapC12 D_5_A, and VapC12 D_94_A proteins were subjected to SDS PAGE and probed with an anti-His antibody.

**S9.**
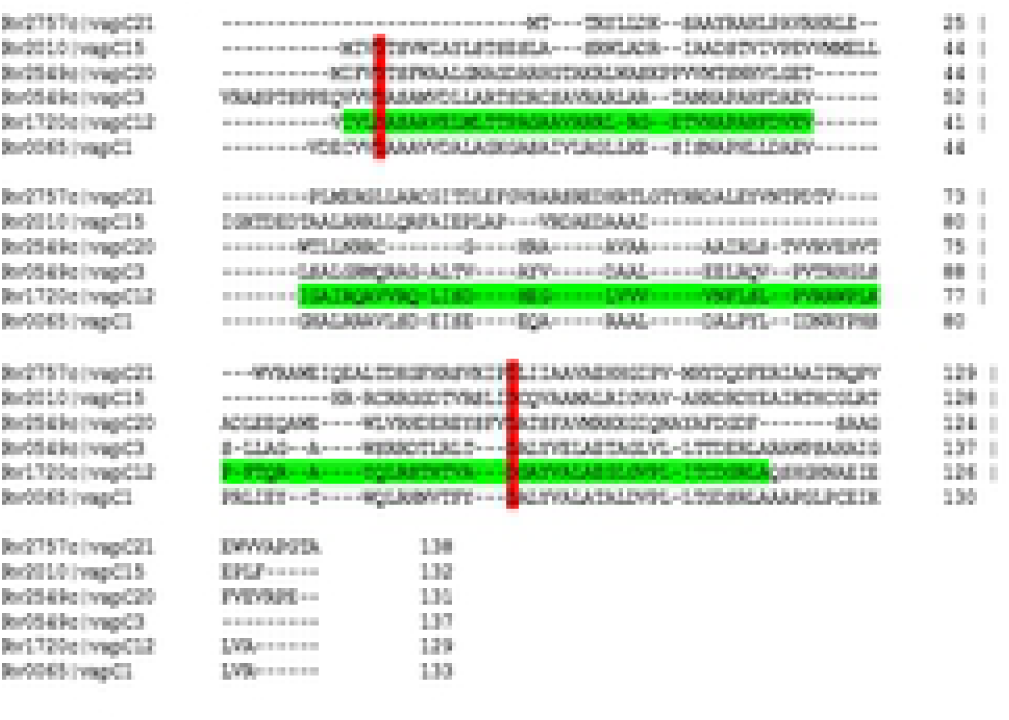
Multiple sequence alignment of VapCs toxins indicating conserved aspartate residues in the PIN domain of toxins.

**S10.**
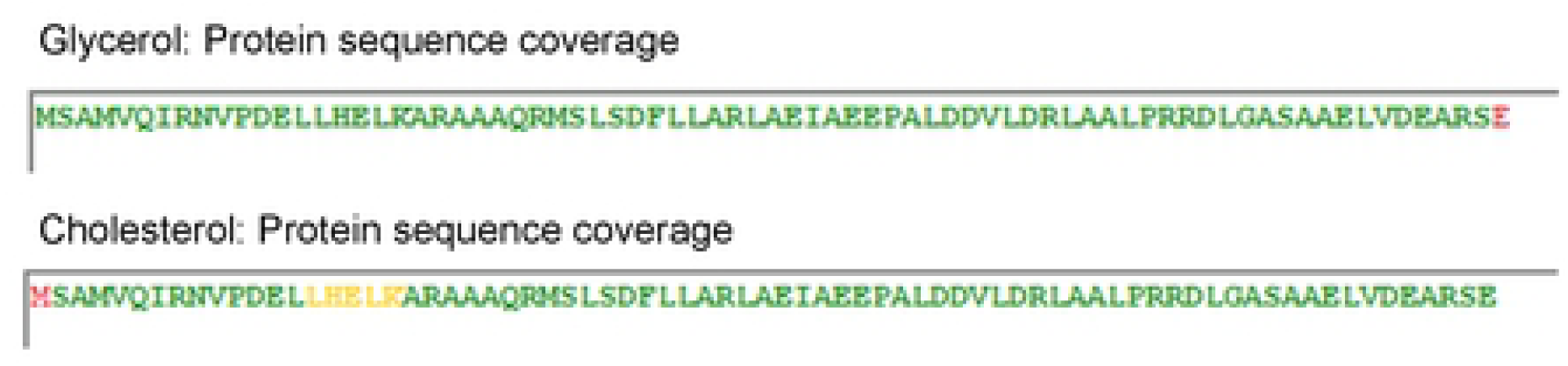
Protein sequence coverage of peptides from BCG:VapBC12 His tagged overexpression strain grown in glycerol and cholesterol media, where grey colour indicates no match or 0 peptide confidence, red colour is >0 and <50 peptide confidence, yellow colour is ≥50 and <95 peptide confidence and green colour is ≥95 peptide confidence.

**S11.**
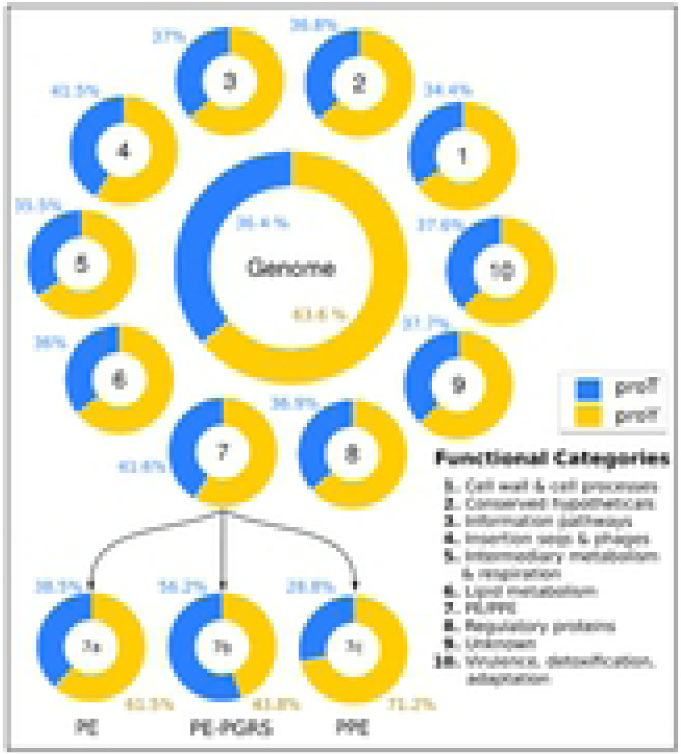
Genome-wide in silico analysis of the codon usage of proT and proY tRNA in each gene belonging to all 10 functional groups in Mtb. Data for codon usage can be obtained from the link provided in material and methods.

**S12.**
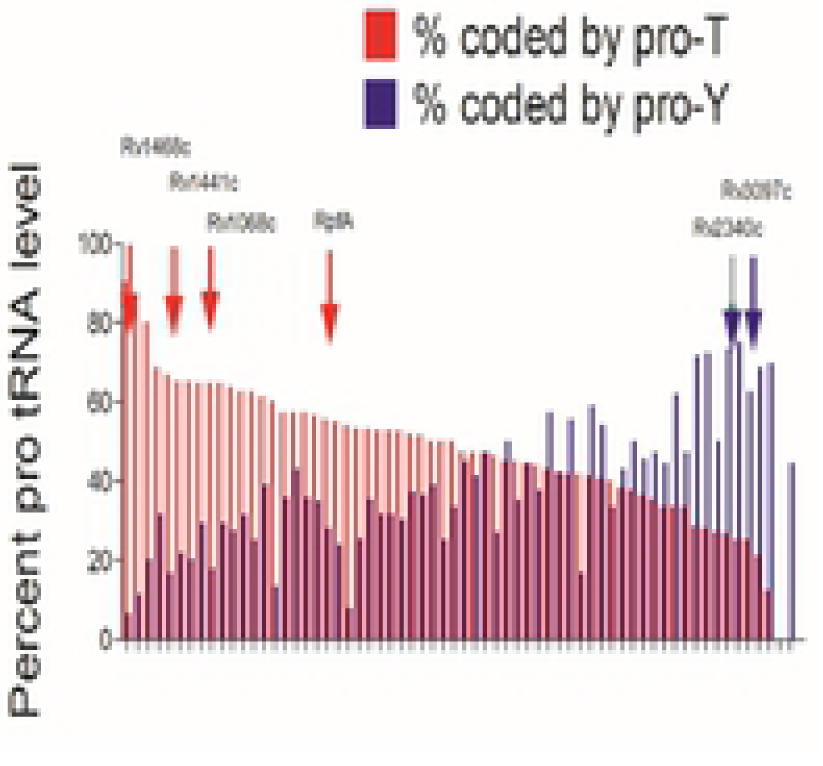
Codon usage of proT and proY tRNA in the PE-PGRS group of genes of Mtb

**S13.**
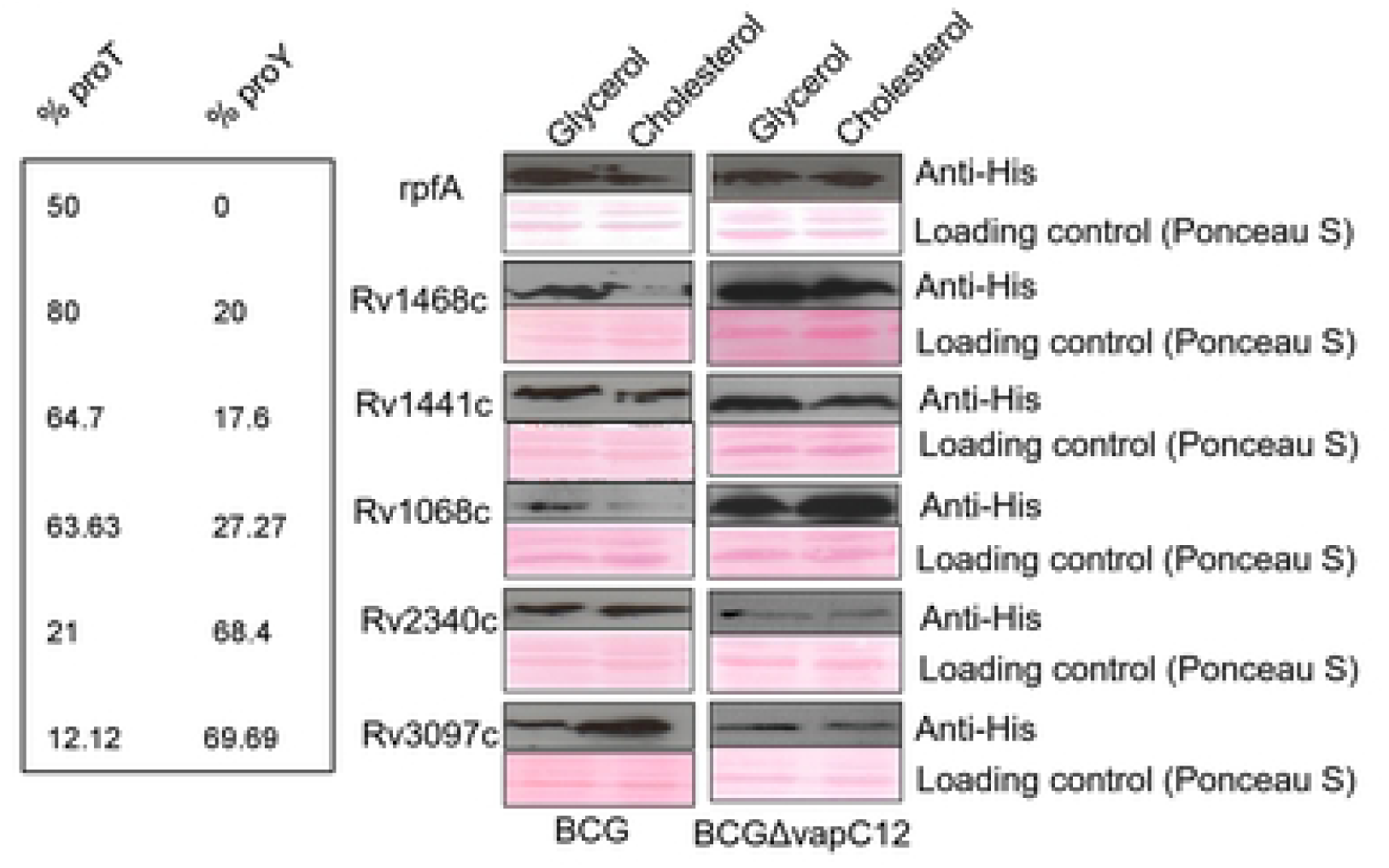
Relative expression of His-tagged PE-PGRS and RpfA proteins in BCG and *vapC12* mutant strains grown in glycerol- and cholesterol-rich media. The protein lysates were prepared from overexpressed strains with an OD of 0.8–1. The samples were run on SDS PAGE and probed with an anti-His antibody.

**S14.**
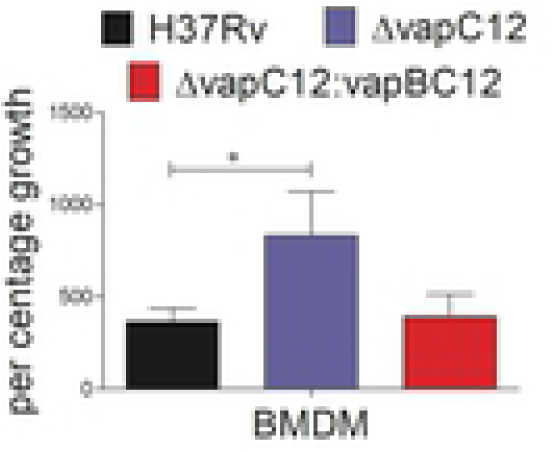
Relative survival of wild-type H37Rv, *ΔvapC12*, and *ΔvapC12:vapBC12* strains in mouse bone marrow-derived macrophages. Infection was performed at MOI of 1, and CFU plating was performed at day 0 and day 7 for bacterial enumeration on 7H11+OADC plates. The experiment was repeated three times, and data plotted represent the mean ± SEM. Data were analysed using unpaired Student’s t test. *P < 0.05 **P < 0.01.

**S15.**
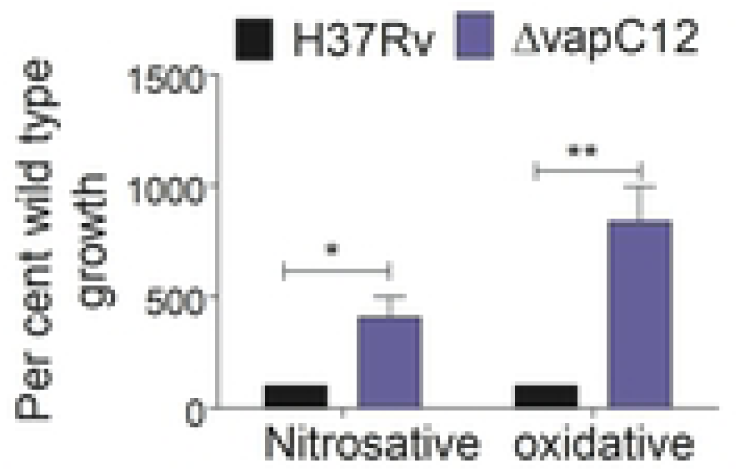
H37Rv and *ΔvapC12* strains were subjected to different stress conditions: nitrosative stress with 200 µM of Deta-NO for 48 hours and oxidative stress with 5 mM of H_2_O_2_ treatment for 6 hours. Percent survival of *ΔvapC12* relative to the wild-type strain was calculated by plating cultures at day zero and respective time points.

**S16.**
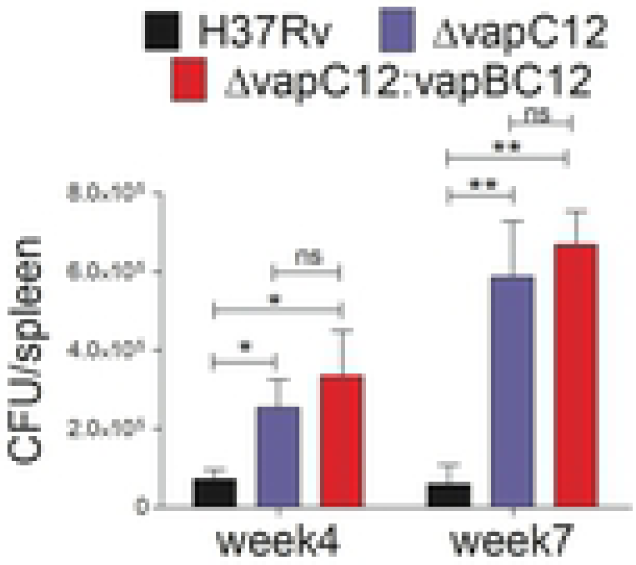
Bacterial load in the spleen of guinea pigs infected with H37Rv, *ΔvapC12*, and *ΔvapC12:vapBC12* strains of Mtb. At designated time points, spleens were homogenized in 4mL of saline, and ten-fold serial dilutions of homogenates were plated on 7H11+OADC plates. Each group constituted six guinea pigs per time point. Data plotted represent the mean ± SEM. Significant differences observed between groups are indicated. Data were analysed using the Mann–Whitney U test with **P < 0.01 and *P < 0.05).

**S17.**
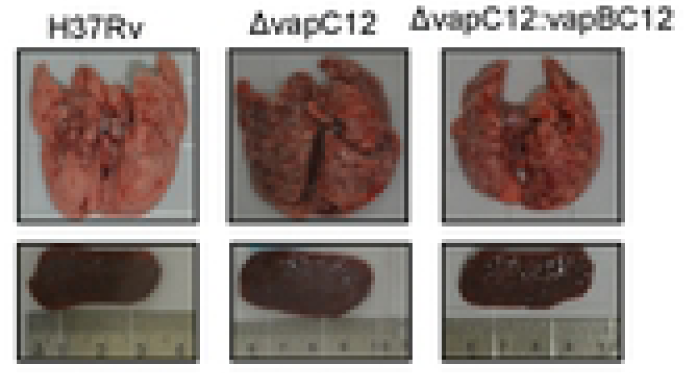
Gross pathology of the lungs and spleen of guinea pigs infected with various strains of Mtb at 7 weeks post infection.

**S18.**
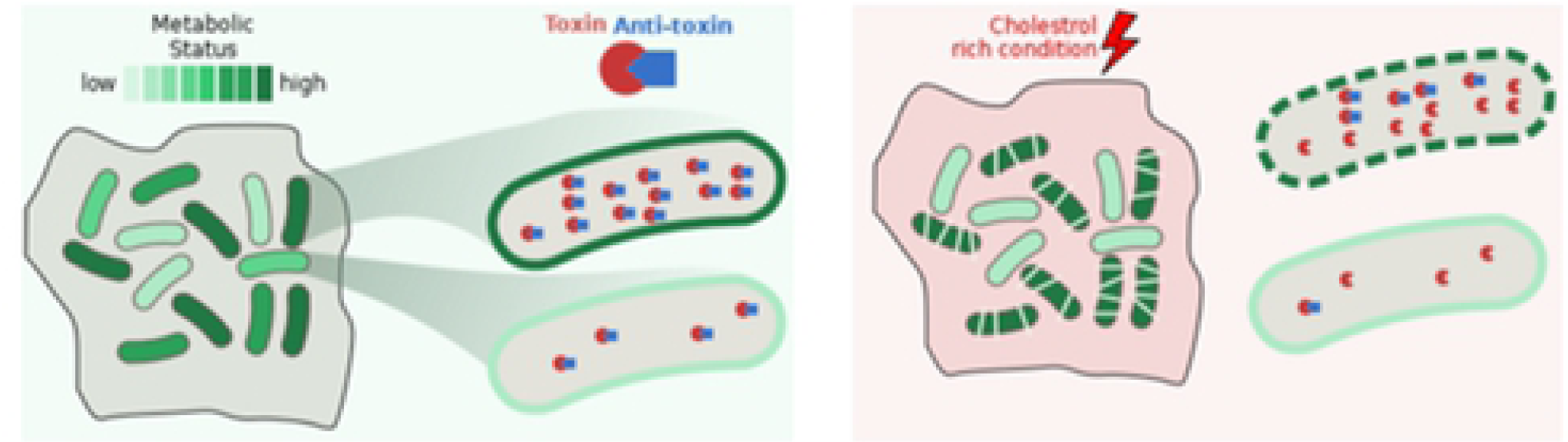
Graphical abstract of the study indicating degradation of AT VapB12 and subsequent activation of toxin VapC12 under cholesterol rich condition. The fast-growing bacteria with higher expression of toxin are eliminated or killed as compared to the slow growing bacteria in the population with less expression of the toxin VapC12.

**Table S1:**
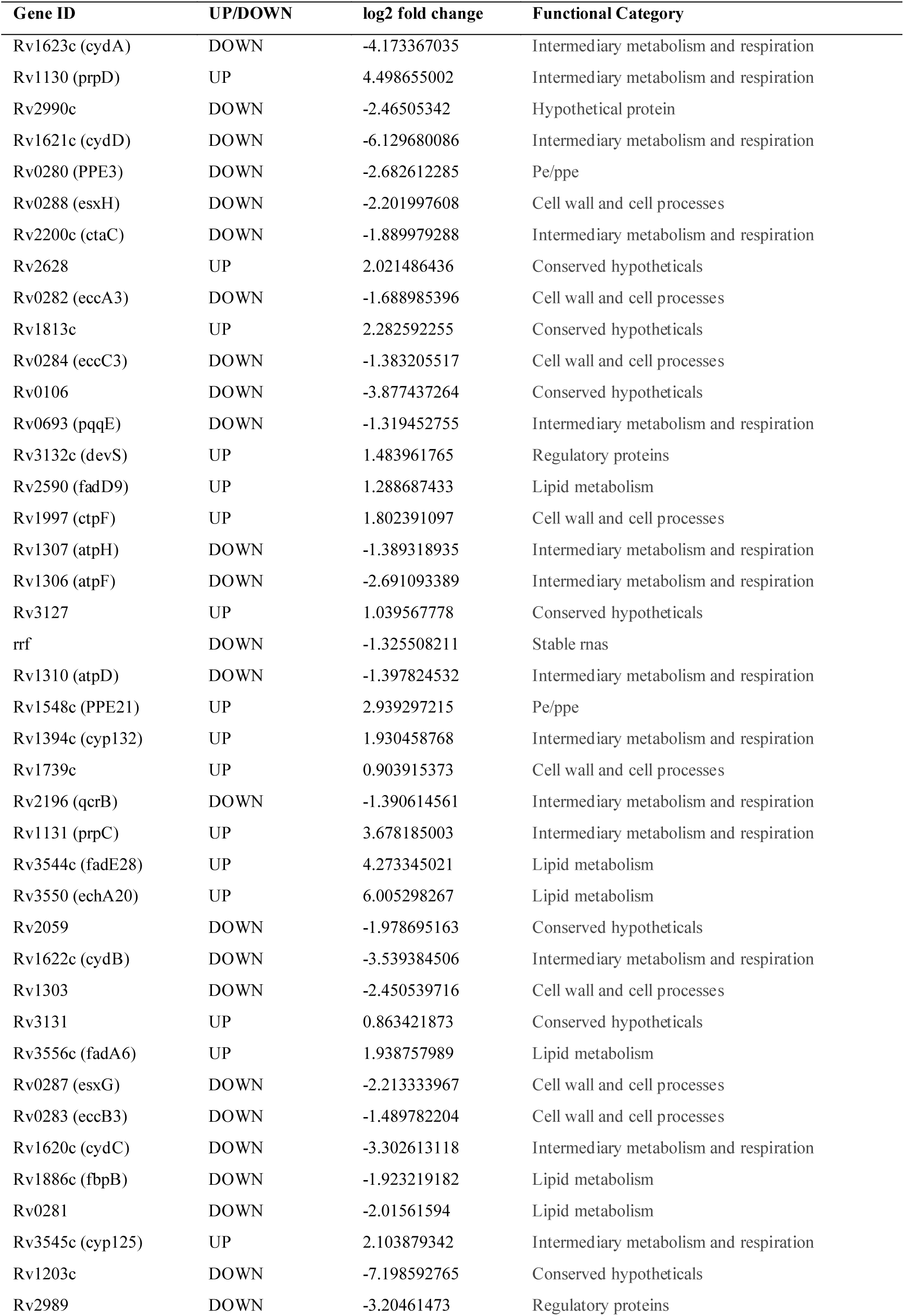
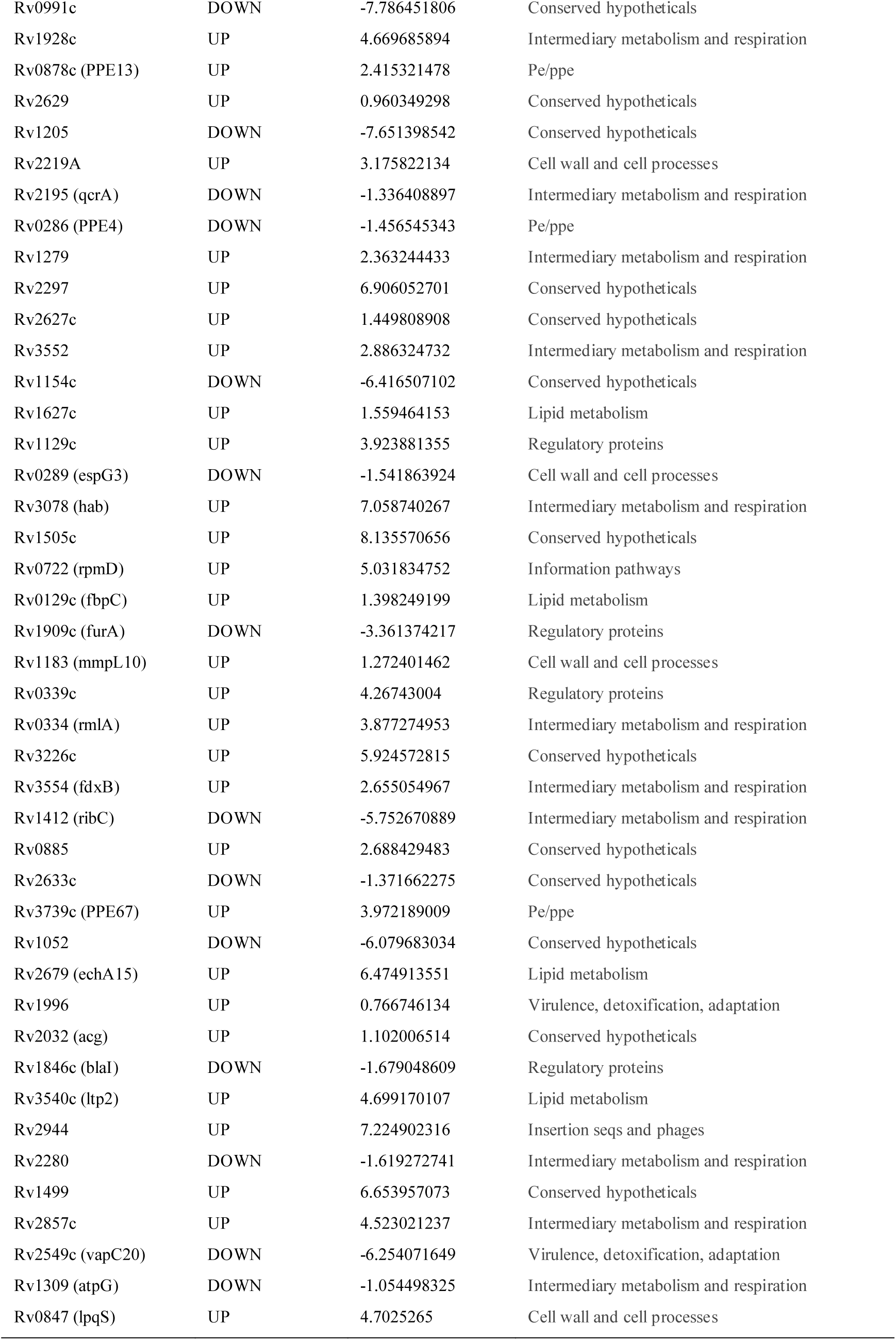
DEGs of H37Rv Cholesterol versus Glycerol.

**Table S2:** DEGs of H37Rv Cholesterol versus ΔvapC12 Cholesterol.

| Gene ID | log2FoldChange | UP/DOWN | Functional Category |
| --- | --- | --- | --- |
| Rv1721c | -3.809774762 | DOWN | Virulence, detoxification and adaptation |
| Rv2378c | -7.834381397 | DOWN | Lipid metabolism |
| Rv1441c | -7.414059049 | DOWN | PE-PPE |
| Rv1740 | -6.647609595 | DOWN | Virulence, detoxification and adaptation |
| Rv2859c | -5.030327611 | DOWN | Intermediary metabolism and respiration |

**Table S3:** Functional categorization of genes with proT codon usage above 60 per cent.

| Gene Id | Functional category | % proT |
| --- | --- | --- |
| Rv0596c | virulence, detoxification, adaptation | 60 |
| Rv1952 | virulence, detoxification, adaptation | 60 |
| Rv2865 | virulence, detoxification, adaptation | 60 |
| Rv2549c | virulence, detoxification, adaptation | 66.66666667 |
| Rv1956 | virulence, detoxification, adaptation | 71.42857143 |
| Rv1982A | virulence, detoxification, adaptation | 75 |
| Rv2863 | virulence, detoxification, adaptation | 80 |
| Rv0300 | virulence, detoxification, adaptation | 100 |
| Rv0550c | virulence, detoxification, adaptation | 100 |
| Rv1103c | virulence, detoxification, adaptation | 100 |
| Rv2760c | virulence, detoxification, adaptation | 100 |
| Rv0737 | regulatory proteins | 60 |
| Rv3557c | regulatory proteins | 60 |
| Rv3050c | regulatory proteins | 62.5 |
| Rv1994c | regulatory proteins | 66.66666667 |
| Rv0348 | regulatory proteins | 71.42857143 |
| Rv2021c | regulatory proteins | 100 |
| Rv3653 | PE/PPE | 60 |
| Rv0872c | PE/PPE | 61.11111111 |
| Rv2853 | PE/PPE | 61.11111111 |
| Rv0354c | PE/PPE | 62.5 |
| Rv2162c | PE/PPE | 62.5 |
| Rv1068c | PE/PPE | 63.63636364 |
| Rv3512 | PE/PPE | 64.28571429 |
| Rv0833 | PE/PPE | 64.70588235 |
| Rv1441c | PE/PPE | 64.70588235 |
| Rv3367 | PE/PPE | 64.70588235 |
| Rv1452c | PE/PPE | 65.2173913 |
| Rv1169c | PE/PPE | 66.66666667 |
| Rv1840c | PE/PPE | 66.66666667 |
| Rv0578c | PE/PPE | 68.29268293 |
| Rv0747 | PE/PPE | 68.42105263 |
| Rv1468c | PE/PPE | 80 |
| Rv3508 | PE/PPE | 85.18518519 |
| Rv3514 | PE/PPE | 90.32258065 |
| Rv0470c | lipid metabolism | 60 |
| Rv0972c | lipid metabolism | 60 |
| Rv2724c | lipid metabolism | 60 |
| Rv2982c | lipid metabolism | 60 |
| Rv3221c | lipid metabolism | 100 |
| Rv0771 | intermediary metabolism and respiration | 60 |
| Rv1851 | intermediary metabolism and respiration | 60 |
| Rv1826 | intermediary metabolism and respiration | 60 |
| Rv2499c | intermediary metabolism and respiration | 60 |
| Rv2511 | intermediary metabolism and respiration | 60 |
| Rv2250A | intermediary metabolism and respiration | 63.63636364 |
| Rv1692 | intermediary metabolism and respiration | 64.70588235 |
| Rv1311 | intermediary metabolism and respiration | 66.66666667 |
| Rv1555 | intermediary metabolism and respiration | 66.66666667 |
| Rv1990A | intermediary metabolism and respiration | 66.66666667 |
| Rv2539c | intermediary metabolism and respiration | 66.66666667 |
| Rv3145 | intermediary metabolism and respiration | 66.66666667 |
| Rv2421c | intermediary metabolism and respiration | 70 |
| Rv3624c | intermediary metabolism and respiration | 70 |
| Rv2754c | intermediary metabolism and respiration | 71.42857143 |
| Rv0558 | intermediary metabolism and respiration | 72.72727273 |
| Rv0814c | intermediary metabolism and respiration | 75 |
| Rv1305 | intermediary metabolism and respiration | 75 |
| Rv3118 | intermediary metabolism and respiration | 75 |
| Rv3154 | intermediary metabolism and respiration | 77.77777778 |
| Rv2537c | intermediary metabolism and respiration | 80 |
| Rv0137c | intermediary metabolism and respiration | 81.81818182 |
| Rv0763c | intermediary metabolism and respiration | 83.33333333 |
| Rv0741 | insertion seqs and phages | 60 |
| Rv1586c | insertion seqs and phages | 60 |
| Rv2014 | insertion seqs and phages | 71.42857143 |
| Rv3638 | insertion seqs and phages | 71.42857143 |
| Rv1702c | insertion seqs and phages | 74.19354839 |
| Rv1584c | insertion seqs and phages | 75 |
| Rv0094c | insertion seqs and phages | 77.27272727 |
| Rv3467 | insertion seqs and phages | 77.27272727 |
| Rv1765A | insertion seqs and phages | 100 |
| Rv1316c | information pathways | 60 |
| Rv2069 | information pathways | 60 |
| Rv2906c | information pathways | 60 |
| Rv2058c | information pathways | 66.66666667 |
| Rv2056c | information pathways | 66.66666667 |
| Rv1643 | information pathways | 66.66666667 |
| Rv0722 | information pathways | 100 |
| Rv2441c | information pathways | 100 |
| Rv3053c | information pathways | 100 |
| Rv3462c | information pathways | 100 |
| Rv1772 | conserved hypotheticals | 60 |
| Rv0678 | conserved hypotheticals | 60 |
| Rv0607 | conserved hypotheticals | 60 |
| Rv3678A | conserved hypotheticals | 60 |
| Rv1590 | conserved hypotheticals | 60 |
| Rv2283 | conserved hypotheticals | 60 |
| Rv2426c | conserved hypotheticals | 60 |
| Rv2438A | conserved hypotheticals | 60 |
| Rv2558 | conserved hypotheticals | 60 |
| Rv3224B | conserved hypotheticals | 60 |
| Rv0323c | conserved hypotheticals | 63.63636364 |
| Rv2257c | conserved hypotheticals | 64.70588235 |
| Rv0078B | conserved hypotheticals | 66.66666667 |
| Rv0181c | conserved hypotheticals | 66.66666667 |
| Rv0530A | conserved hypotheticals | 66.66666667 |
| Rv1120c | conserved hypotheticals | 66.66666667 |
| Rv2820c | conserved hypotheticals | 66.66666667 |
| Rv2239c | conserved hypotheticals | 66.66666667 |
| Rv2342 | conserved hypotheticals | 66.66666667 |
| Rv2923c | conserved hypotheticals | 66.66666667 |
| Rv3472 | conserved hypotheticals | 66.66666667 |
| Rv3033 | conserved hypotheticals | 71.42857143 |
| Rv0028 | conserved hypotheticals | 75 |
| Rv1890c | conserved hypotheticals | 75 |
| Rv2603c | conserved hypotheticals | 75 |
| Rv1066 | conserved hypotheticals | 80 |
| Rv2049c | conserved hypotheticals | 100 |
| Rv0378 | conserved hypotheticals | 100 |
| Rv1893 | conserved hypotheticals | 100 |
| Rv1993c | conserved hypotheticals | 100 |
| Rv2548A | conserved hypotheticals | 100 |
| Rv2738c | conserved hypotheticals | 100 |
| Rv3440c | conserved hypotheticals | 100 |
| Rv0011c | cell wall and cell processes | 60 |
| Rv0431 | cell wall and cell processes | 60 |
| Rv1463 | cell wall and cell processes | 60 |
| Rv1973 | cell wall and cell processes | 60 |
| Rv2856 | cell wall and cell processes | 60 |
| Rv2936 | cell wall and cell processes | 61.53846154 |
| Rv3864 | cell wall and cell processes | 61.9047619 |
| Rv3312A | cell wall and cell processes | 62.5 |
| Rv0583c | cell wall and cell processes | 65 |
| Rv2732c | cell wall and cell processes | 66.66666667 |
| Rv3277 | cell wall and cell processes | 66.66666667 |
| Rv0476 | cell wall and cell processes | 75 |
| Rv1881c | cell wall and cell processes | 75 |
| Rv2301 | cell wall and cell processes | 75 |
| Rv3271c | cell wall and cell processes | 75 |
| Rv0900 | cell wall and cell processes | 100 |
| Rv0039c | cell wall and cell processes | 100 |
| Rv0288 | cell wall and cell processes | 100 |
| Rv2520c | cell wall and cell processes | 100 |
| Rv3789 | cell wall and cell processes | 100 |
| Rv3857c | cell wall and cell processes | 100 |

**Table S4:**
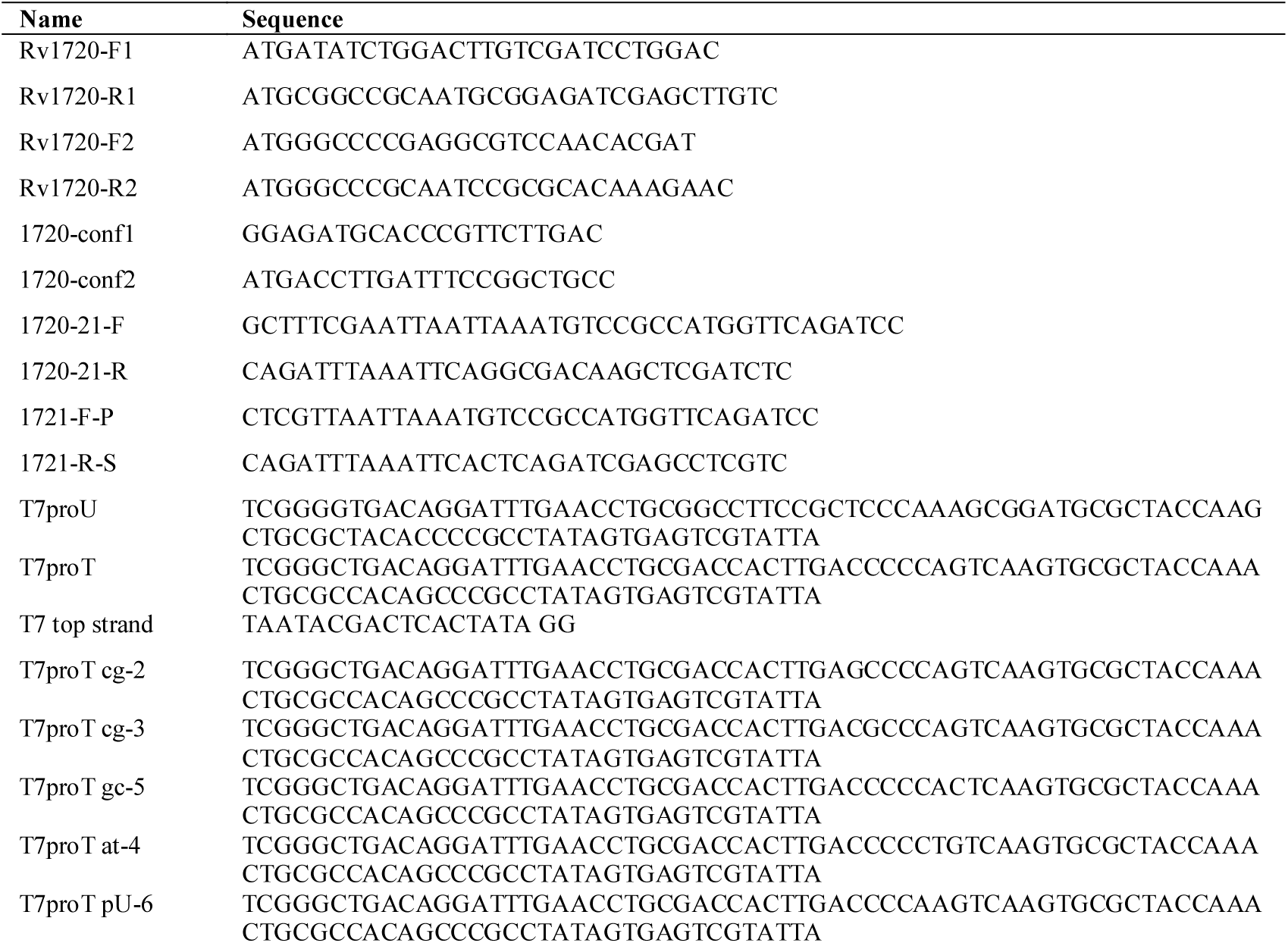
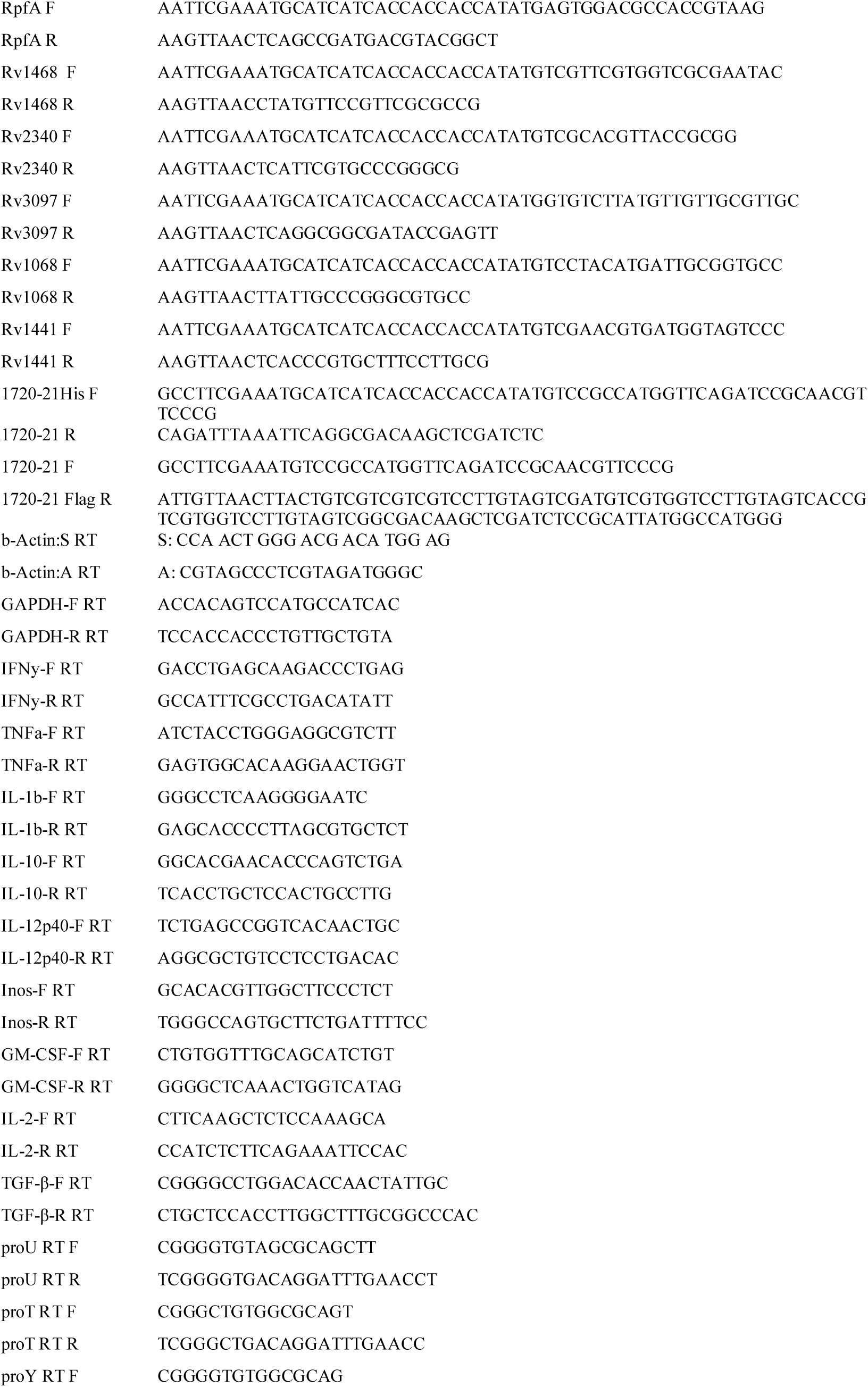
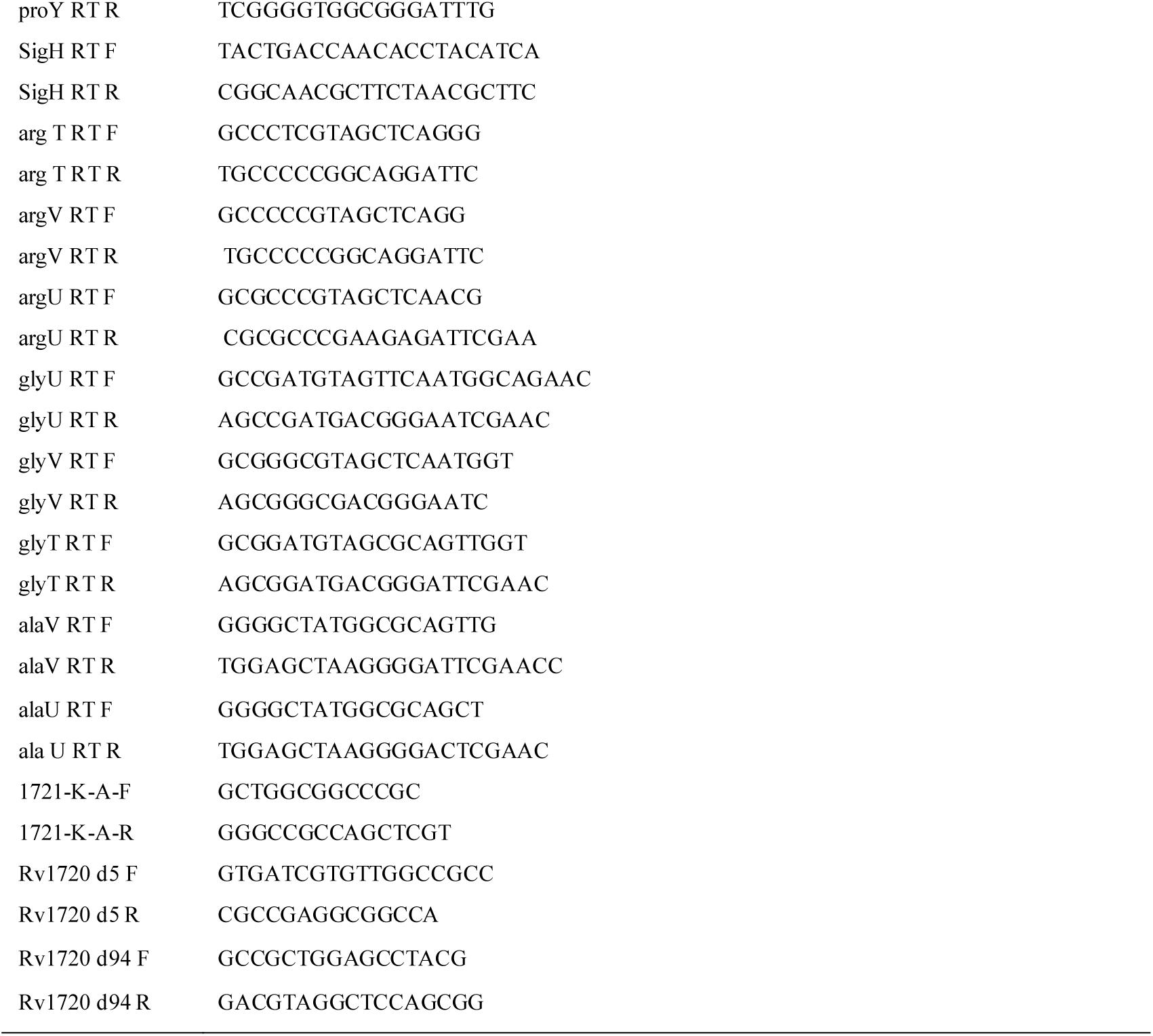
Primers used in the study.

**Table S5:**
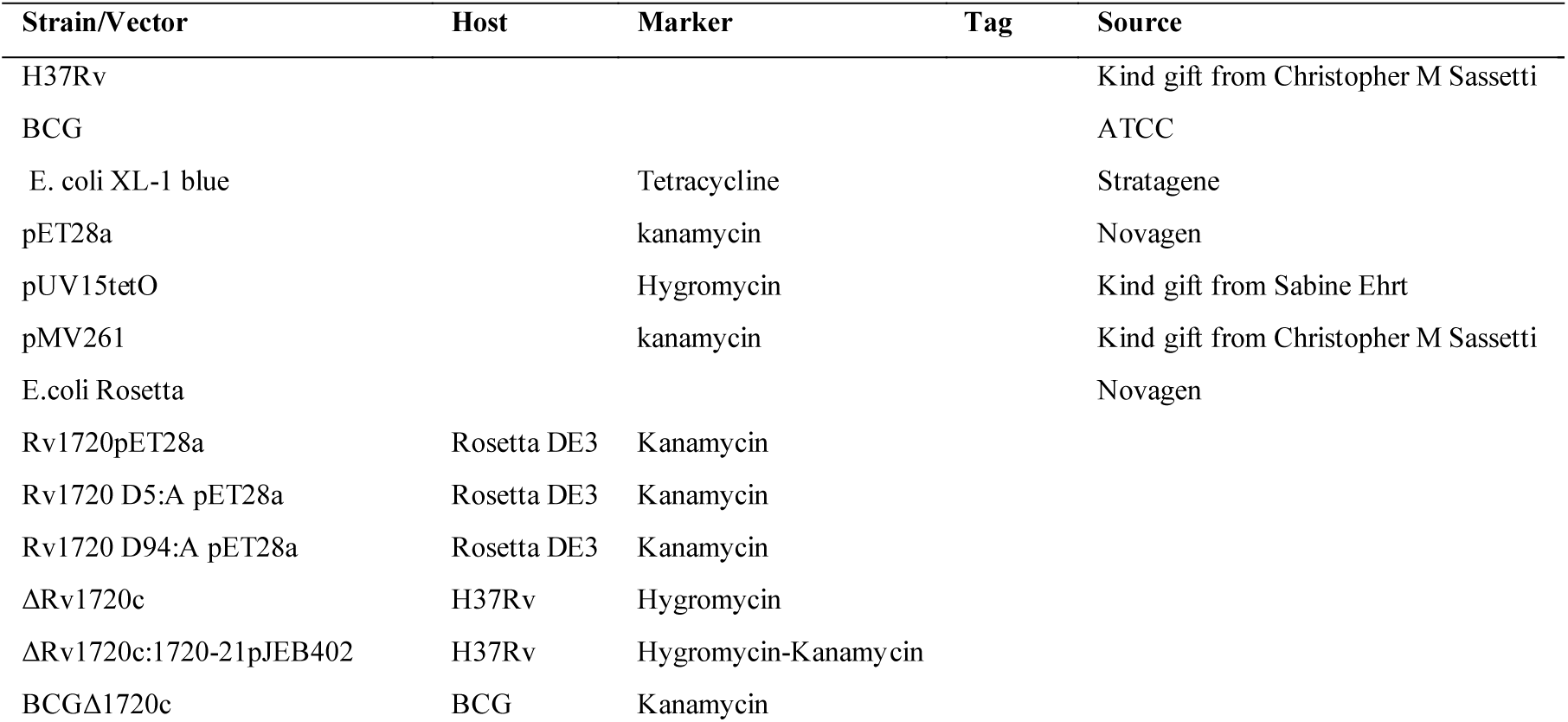
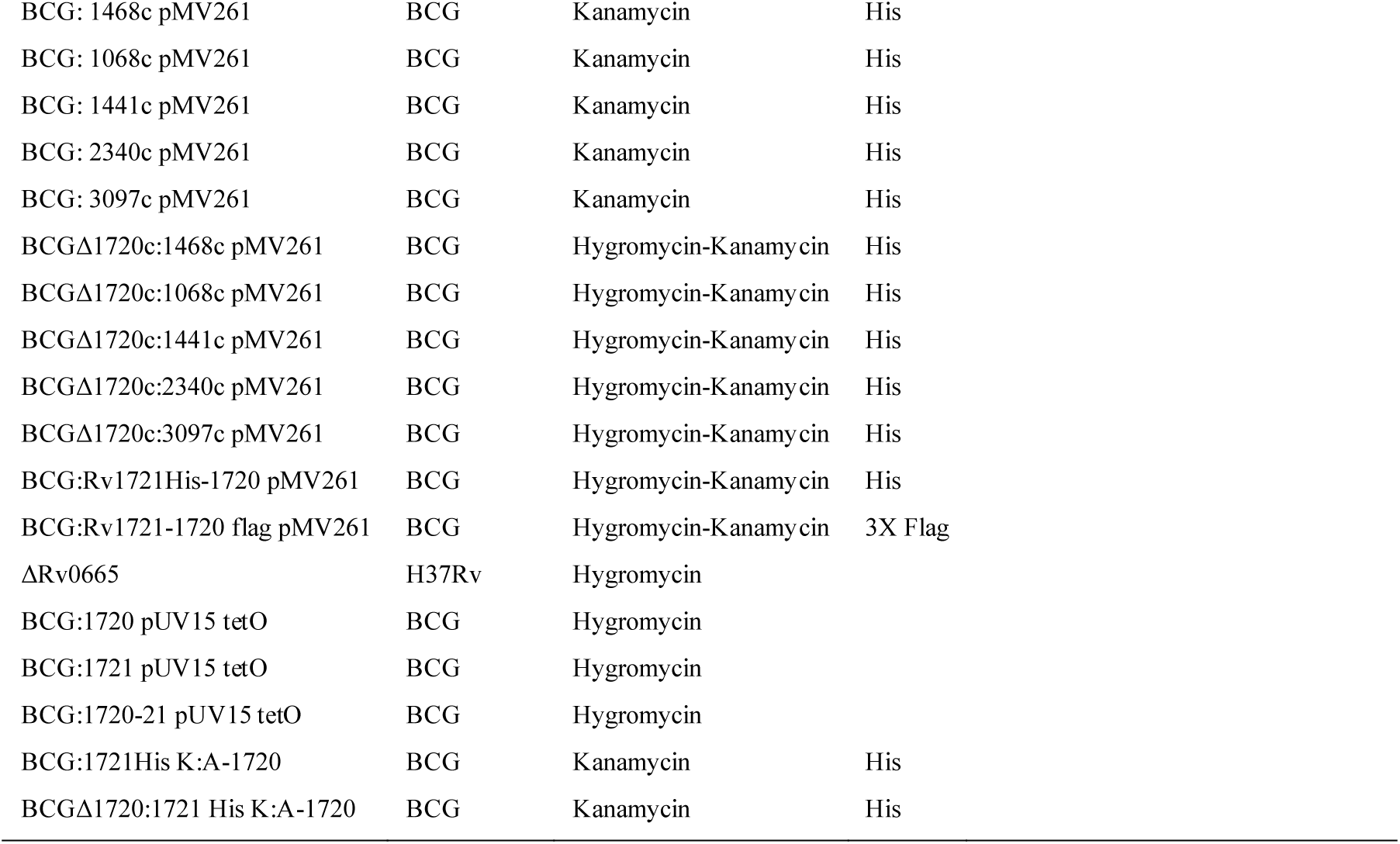
Strains used in this study.

